# X Chromosome Dosage Modulates Multiple Molecular and Cellular Properties of Mouse Pluripotent Stem Cells Independently of Global DNA Methylation Levels

**DOI:** 10.1101/291450

**Authors:** Juan Song, Adrian Janiszewski, Natalie De Geest, Lotte Vanheer, Irene Talon, Mouna El Bakkali, Taeho Oh, Vincent Pasque

**Affiliations:** KU Leuven - University of Leuven, Department of Development and Regeneration, Herestraat 49, B-3000 Leuven, Belgium

## Abstract

During early mammalian development, the two X-chromosomes in female cells are active. Dosage compensation between XX female and XY male cells is then achieved by X-chromosome inactivation in female cells. Reprogramming female mouse somatic cells into induced pluripotent stem cells (iPSCs) leads to X-chromosome reactivation. The extent to which increased X-chromosome dosage (X-dosage) in female iPSCs leads to differences in the molecular and cellular properties of XX and XY iPSCs is still unclear. We show that chromatin accessibility in mouse iPSCs is modulated by X-dosage. Specific sets of transcriptional regulator motifs are enriched in chromatin with increased accessibility in XX or XY iPSCs. We show that the transcriptome, growth and pluripotency exit are also modulated by X-dosage in iPSCs. To understand the mechanisms by which increased X-dosage modulates the molecular and cellular properties of mouse pluripotent stem cells, we used heterozygous deletions of the X-linked gene *Dusp9* in XX embryonic stem cells. We show that X-dosage regulates the transcriptome, open chromatin landscape, growth and pluripotency exit largely independently of global DNA methylation. Our results uncover new insights into X-dosage in pluripotent stem cells, providing principles of how gene dosage modulates the epigenetic and genetic mechanisms regulating cell identity.

## INTRODUCTION

Pluripotent stem cells (PSCs) are important for modeling development and diseases and for the design of future regenerative medicine approaches [1]. A key question in the field is what mechanisms underlie the establishment and maintenance of pluripotency. Somatic cells can be reprogrammed into induced PSCs (iPSCs) by transcription factor (TF) overexpression [2], and mouse embryonic stem cells (ESCs) can be derived directly from early embryos [3]. Both cell types have the capacity to self-renew and maintain embryonic lineage differentiation potential in culture [4]. It is of outstanding interest to understand what epigenetic and genetic mechanisms influence the molecular and functional properties of PSCs.

Several mammalian species including mice and human have adopted X-chromosome inactivation as a means to compensate between the genetic imbalance of XX female and XY male cells (reviewed in [5]). X-chromosome inactivation is established during early embryogenesis following the expression of the long non-coding RNA *Xist*, and maintained in most somatic cells. Female cells undergo X-chromosome reactivation in the mouse inner cell mass (ICM) resulting in two active X-chromosomes (XaXa), a state maintained in female ESCs [6,7]. X-chromosome reactivation is also induced following somatic cell reprogramming to iPSCs [7] (reviewed in [8]). XaXa is a hallmark of mouse naive pluripotency, the latter is characterized by unbiased embryonic lineage differentiation potential. Consequently, XX mouse ESCs have a higher dose of X-linked gene transcripts and hence an increased X-to-autosome gene expression ratio compared with XY cells. Increasing evidence suggests that the presence of two active X-chromosomes can modulate the molecular and functional properties of mammalian PSCs [9–20]. Work over the past decade showed that XX female ESCs exhibit global DNA hypomethylation affecting most genomic features including imprint control regions [13–21]. Recent work showed that XX female iPSCs also display global hypomethylation [22,23]. Differences in global DNA methylation have been attributed to X-dosage since female XO cells display male-like DNA methylation levels [13,15,17]. Thus, mouse ESCs and iPSCs both show global DNA methylation differences due to X-dosage.

It was also discovered that XX female ESCs show increased expression of several pluripotency-associated transcripts, and display delayed pluripotency exit, suggesting that features of naive pluripotency are promoted in XX female ESCs [15,24]. Differences in transcription have also been attributed to X-dosage since XO female ESCs, or *Xist*-induced X-chromosome inactivation, are associated with male-like pluripotency-associated gene expression and pluripotency exit [15]. Despite the potential influence of X-dosage on iPSCs, X-dosage has been largely ignored in iPSC reprogramming studies so far, and it remains unclear if X-dosage influences the molecular features of iPSCs beyond DNA methylation. Therefore, it is important to determine the potential influence of X-dosage on the molecular and cellular properties of iPSCs, which could influence mechanistic studies of reprogramming. Despite its importance, a systematic comparison of transcriptional states, open chromatin landscapes, growth and pluripotency exit in XX female and XY male mouse iPSCs has not yet been performed.

While several advances have been made, the molecular pathways by which XaXa modulate pluripotency remain incompletely understood [10]. At the mechanistic level, two active X-chromosomes inhibit MAPK and GSK3 signaling [12,15], and global DNA hypomethylation has been attributed to reduced expression of DNMT3A and DNMT3B [13], or DNMT3L [14], or UHRF1 [17,19,22] in XX female ESCs/iPSCs. More recently, it was discovered that increased dosage of the X-linked MAPK inhibitor *Dusp9* (dual-specificity phosphatase 9) is in part responsible for inhibiting DNMT3A/B/L and global DNA methylation in XX female ESCs [17]. The expression level of *Dusp9* is higher in XX ESCs than in XY ESCs and overexpression of *Dusp9* in XY male ESCs induced female-like global DNA hypomethylation and a female-like proteome [17]. Conversely, heterozygous deletion of *Dusp9* in XX female ESCs restored male-like global DNA methylation, suggesting that *Dusp9* is responsible for MAPK-mediated DNMT3A/B repression in XX female ESCs. However, whether *Dusp9* heterozygous deletion in XX female ESCs has effects on the transcriptional regulatory network, open chromatin landscape and pluripotency exit has not yet been explored. In addition, how and which X-linked genes modulate the pluripotency gene network of naive PSCs remains unclear [10]. Furthermore, novel insights may be gained by identification of cis-regulatory elements that drive X-dosage-specific pluripotent stem cell states.

Here, in order to investigate the influence of X-dosage on iPSCs, we systematically compared multiple molecular and cellular properties of mouse XX female and XY male iPSCs at different passages. We found that X-dosage is associated with differences in chromatin accessibility, cell growth, the transcriptome and pluripotency exit in early passage iPSCs, which are subsequently resolved as a result of X-chromosome loss in female iPSCs upon prolonged culture. We further investigated the regulatory landscape of XX female and XY male iPSCs and ESCs. We found that thousands of chromatin regions differ in accessibility between XX and in XY iPSCs. Motif discovery analysis identified that chromatin more accessible in XX female iPSCs is enriched for binding sites of key pluripotency regulators including KLF/ZIC3/NANOG, suggesting stabilization of the naive pluripotency regulatory network via these regulators. By contrast, chromatin sites more accessible in XY iPSCs are enriched for AP-1 motifs, downstream effectors of signaling pathways including MAPK. We show that XY iPSCs grow faster than XX iPSCs, irrespective of culture conditions. We further demonstrate that *Dusp9* heterozygous XX female ESCs maintain female-like chromatin accessibility, growth, and delayed exit from pluripotency in the presence of male-like global DNA methylation. Altogether, our study uncovers X-dosage as a previously unrecognized modulator of chromatin accessibility and of growth in pluripotent stem cells. Furthermore, we redefine the effects of X-dosage on the pluripotency transcriptome, revealing the uncoupling of DNA methylation from chromatin accessibility. This provides principles of using gene dosage in designing experiments to understand the epigenetic and genetic mechanisms regulating cell identity.

## RESULTS

### Differences in Transcriptional Landscapes and Pluripotency Exit Correlate with the Presence of Two Active X-chromosomes in iPSCs

To explore the importance of X-dosage on the transcriptome and pluripotency exit of mouse iPSCs, we derived XX female and XY male iPSC lines. We used isogenic mouse embryonic fibroblasts (MEFs) carrying a tetO inducible transgene encoding the reprogramming factors *Oct4, Sox2, Klf4* and *c-Myc* in the *Col1A* locus and the reverse tetracycline transactivator (M2rtTA) in the *Rosa26* locus (Fig 1A, S1 Table) [23,25]. After 16 days of doxycycline (dox) treatment to induce reprogramming, 10 female and 11 male iPSC lines were expanded on feeders in the presence of serum and LIF (S/L) in the absence of dox (Fig 1A), or adapted to dual ERK/GSK3 inhibition and LIF conditions (2i/L). This scheme allowed us to directly compare female and male iPSCs without the influence of differences in genetic background, reprogramming system or derivation method. Both female and male iPSCs could be propagated over multiple passages while maintaining their morphology, indicative of self-renewal, and expressed pluripotency-associated factors NANOG and DPPA4 (Fig 1B/C, S1 Fig A/B). Thus, derivation of isogenic female and male iPSCs allowed us to systematic compare the transcriptome and epigenome of these cells.

**Fig 1.**
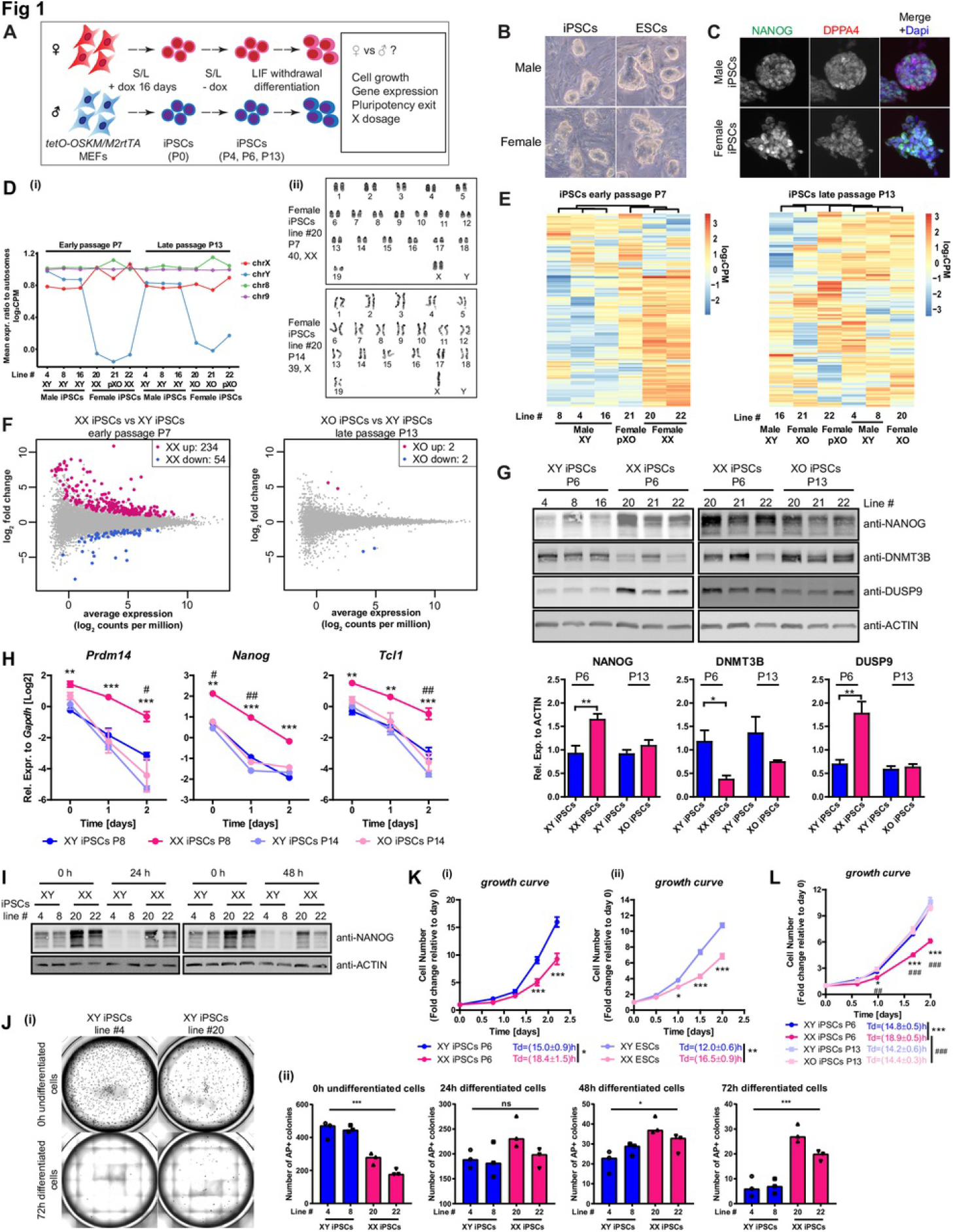
Two X-chromosomes Modulate the Transcriptome, Cellular Growth and Pluripotency Exit in Mouse iPSCs. (A) Scheme of female and male iPSCs derivation, characterization and differentiation.
(B) Representative images of female and male iPSCs and ESCs grown on feeders in S/L condition.
(C) Immunofluorescence analysis for NANOG and DPPA4 in iPSCs grown in S/L. Representative images of all lines examined for NANOG (Red), DPPA4 (Green) and Dapi (Blue, nuclei counterstaining) are shown.
(D) Mean expression ratio to autosomes for sex chromosomes and chromosomes 8 and 9. The dosage of X and Y linked genes was used to infer XX, XY, XO and partial XO (pXO) genotypes.
(E) Unsupervised hierarchical clustering of top 200 most variable autosomal genes in XY, XX, pXO and XO iPSCs. Early passage iPSCs cluster by X-dosage, late passage iPSCs do not.
(F) DEG analysis, identifying clear differences between XX female and XY male iPSCs, but not XO female and XY male iPSCs.
(G) Western blot analysis for NANOG, DNMT3B and DUSP9 protein in iPSCs grown in S/L in both early and late passages. Lower panel: quantification using ACTIN as a loading control. The protein values are represented as averages (±SEM) of male and female iPSC lines (n=3 lines each) in early passage (P6) and late passage (P13), respectively. Statistical significance was analysed using the unpaired, two-tailed t-test (*p<0.05, **p<0.01).
(H) RT-qPCR analysis for pluripotency-associated gene expression during LIF withdrawal differentiation of both early passage and late passage iPSCs. Results are presented as averages (±SEM) of three XY male and three XX female iPSC lines in early passage (P8) and the same lines in late passage (P14), respectively. **p<0.01, ***p<0.001, P8 XX iPSCs vs P8 XY iPSCs. #p<0.05, ##p<0.01, P14 XO iPSCs vs P14 XY iPSCs. Two-way repeated-measures ANOVA with Bonferroni posttests.
(I) Western blot analysis for NANOG during pluripotency exit. The time after LIF withdrawal is indicated.
(J) XX and XY iPSCs were subject to 0h, 24h, 48h and 72h of LIF withdrawal before replating 5000 cells/well on feeders in 12-well plates. (i) Representative images of AP+ colonies for replated XX and XY iPSCs after 5 days in 2i/L. (ii) The number of AP+ colonies after 5 days in 2i/L is indicated. Statistical significance was analysed using the one-way ANOVA with Tukey’s multiple comparisons test (*p<0.05, ***p<0.0001).
(K) Growth curves and doubling times of XX and XY iPSCs (i) and ESCs (ii) in S/L condition. Cells were counted at the indicated time points and presented as fold changes relative to averages (±SEM) of XY and XX iPSC lines at 0h (three lines each, n=1, left panel) in early passage (P6) and of XY and XX ESCs (one line each, n=3, right panel), respectively. Growth curve: *p<0.05, ***p<0.001, XY iPSCs vs XX iPSCs, XY ESCs vs XX ESCs, two-way repeated-measures ANOVA with Bonferroni posttests. Doubling time (Td): *p<0.05, **p<0.01, XY iPSCs vs XX iPSCs, XY ESCs vs XX ESCs, unpaired two-tailed t-test.
(L) As in (K) but for XY, XX and XO iPSCs. Growth curve: *p<0.05, ***p<0.001, P6 XX iPSCs vs P6 XY iPSCs; ##p<0.01, ###p<0.001, P13 XX iPSCs vs P13 XY iPSCs; two-way repeated-measures ANOVA with Bonferroni posttests. Td: ***p<0.01, P6 XX iPSCs vs P6 XY iPSCs; ###p<0.001, P13 XO iPSCs vs P13 XY iPSCs; unpaired two-tailed t-test.

First, we confirmed that XX female iPSCs reactivated the inactive X-chromosome, a hallmark of naive pluripotency [26], using RNA-seq analysis (Fig 1D). These results were also in agreement with an independent single-cell level assay using RNA *in situ*-hybridization for X-linked gene *Tsix* (S1 Fig C and [27]). XX female ESCs are prone to lose one of the two active X-chromosomes upon extended *in vitro* cell culture [13,14,17–19,28,29], and we recently showed that early passage XX female iPSCs are XaXa and become XO iPSCs upon passage [23]. To infer X-chromosome loss in our iPSC lines, we measured the average X-chromosome-to-autosome gene expression ratio using RNA-seq (Fig 1D). We found that early passage XX female iPSCs had increased X-dosage, in agreement with the XaXa state of female iPSCs [30]. However, female iPSCs at late passage showed reduced X-dosage, consistent with X-chromosome loss, and we termed these cells XO iPSCs. In addition, we found that X-chromosome loss in female iPSC lines displayed clonal variability. One early passage and one late passage female iPSC line showed partial X-dosage, consistent with partial X loss, which we termed partial XO (pXO) iPSCs. In further support of our finding that XX female iPSCs undergo X-chromosome loss rather than X-chromosome inactivation, we designed a simple qPCR assay to determine the X/autosome genomic DNA ratio by measuring four X-linked genes (*Tfe3*, *Bcor*, *Pdha1* and *Mid1*, located on either distal region on the X-chromosome) and the autosomal gene *Gapdh*. We confirmed that late passage iPSCs were XO (S1 Fig D). Karyotype analyses corroborated these results (Fig 1D(ii)). These observations are consistent with X-chromosome reactivation during reprogramming followed by X-chromosome loss in female iPSCs.

Using RNA-seq of XX, XY and XO iPSCs grown in S/L, we asked whether the transcriptome of iPSCs is influenced by X-dosage. Unsupervised clustering of the top 200 most variable autosomal genes, or genes associated with stem cell maintenance, distinguished early passage XX female and XY male iPSCs (Fig 1E, S1 Fig E). However, XO female and XY male late passage iPSCs could not be distinguished, indicating that X-dosage rather than sex modulates the transcriptome of iPSCs. Furthermore, gene expression analysis identified 288 differentially expressed genes (DEGs) between XX and XY iPSCs, whereas there were only 4 DEGs between XO female and XY male iPSCs (Fig 1F, 1.5 fold, FDR=0.05, S2 Table). Using reverse transcription (RT) quantitative real-time PCR (qPCR) we found that in S/L, XX female iPSC lines consistently expressed higher levels of pluripotency-associated genes *Prdm14*, *Nanog* and *Tcl1* compared with XY male iPSCs (S1 Fig F). Western blot analysis showed that XX female iPSCs had increased NANOG protein levels compared with XY male and XO female iPSCs (Fig 1G). These marked differences between XX and XY iPSCs are consistent with patterns observed in mouse ESCs [15,17] (S1 Fig G/H, S3 Table), in agreement with the notion that iPSCs are molecularly equivalent to ESCs. Yet, X-dosage has been largely ignored in mechanistic iPSC reprogramming studies so far. Importantly, differences between XX and XY iPSCs cannot be attributed to differences in genetic background since these differences were found when comparing cells of the same genetic background. Thus, reprogramming to iPSCs results in differences in the transcriptome of iPSCs, some of which can be attributed to differences in X-dosage.

Next, we investigated the extent to which X-dosage affects exit from pluripotency in iPSCs. We subjected XX female, XY male and XO female iPSCs to LIF withdrawal-mediated differentiation and measured the downregulation of pluripotency-associated genes by RT-qPCR (Fig 1A/H). Exit from pluripotency was delayed in XX iPSCs for *Prdm14*, *Nanog* and *Tcl1*, but not for XY and XO iPSCs (Fig 1H). We confirmed these results using an alternative differentiation protocol that mimics epiblast differentiation (S1 Fig I/J) [15,31]), and also at the protein level (Fig 1I). These differences had functional consequences on pluripotency exit: replating an equal number of XX or XY cells before and after LIF withdrawal followed by 2i/L culture confirmed delayed pluripotency exit in XX cells (Fig 1J). Thus, XX female iPSCs functionally exit pluripotency with delayed kinetics compared with XY male and XO female iPSCs. Altogether, these findings show that early passage iPSCs display previously unrecognized X-dosage specific behavior in transcriptome, including pluripotency gene expression, and in pluripotency exit kinetics, consistent with X-dosage differences in ESCs [15,24,32].

### X-Dosage Modulates Cellular Growth in Mouse iPSCs and ESCs

To determine the effect of X-dosage on cell growth, we counted the number of XX female and XY male iPSCs over two days starting from the same amount of cells. We found that XX female iPSC lines grew slower than XY male iPSCs, with a doubling time (Td) extended by ^∼^3.4 hours compared with XY male iPSCs grown in S/L (Td XX female iPSCs=18.4.±1.5hr vs Td XY male iPSCs=15.0±0.9hr) (Fig 1K). XX female ESCs also grew slower than XY male ESCs (Fig 1K). The delayed growth of XX female iPSCs was attributed to the presence of two active X-chromosomes, since XO female iPSCs behaved like XY male iPSCs (Fig 1L). The differences in growth of XX female and XY male iPSCs and ESCs did not depend on culture conditions because XX female ESCs and iPSCs still grew slower than XY male cells in 2i/LIF (S1 Fig K). XX female mouse and human embryos show a delay in post-implantation development that has been attributed to the presence of two X-chromosomes in female cells [32]. Our observations support the idea that the growth delay of XX female mammalian embryos is recapitulated *in vitro* in iPSCs and ESCs cultures, providing a new platform to study this process [32].

To assess the effect of X-dosage on the cell cycle, we used 5-ethynyl-2’-deoxyuridine (EdU) incorporation and propidium iodide (PI) staining in combination with flow cytometry to determine the distribution of cells over the different phases of the cell cycle. We found that the majority of both XX and XY iPSCs and ESCs reside in S-phase, in line with the literature. The proportion of XX female iPSCs and ESCs in S-phase was larger than that of XY male iPSCs and ESCs, whereas the number of XX female iPSCs and ESCs in the G1-phase was smaller than that of XY cells (S1 Fig L(i)). To further validate these results, we introduced a Fluorescence Ubiquitination Cell Cycle Indicator (FUCCI) into XX female and XY male ESCs. This system provides for direct fluorescent visualization of ESCs in G1 phase, G1/S transition, or S/G2/M phase [33]. This analysis confirmed, for XX ESCs, an increase in the proportion of cells in S phase, and a reduced proportion of XX cells in G1 phase, compared with XY ESCs (S1 Fig L(ii)).

What might be the functional relevance of differences in cell growth between cells with one or two active X-chromosomes? It has been suggested that the presence of two X-chromosomes slows down development to ensure that the cells progress through X-chromosome inactivation [15]. We sought to test, *in vitro*, the hypothesis that reduced X-dosage provides a competitive growth advantage to cells that have undergone X-chromosome inactivation. We mixed XX female ESCs and GFP-labeled XY male ESCs in different ratios and followed the proportion of labeled cells over time. We found that the increased cell growth of XY male ESCs can provide a small advantage over a 8 day period (S1 Fig M). Collectively, these observations support the idea that cell growth is decreased as a result of increased X-dosage in pluripotent cells *in vitro* and *in vivo*.

### Influence of X-Dosage on Chromatin Accessibility in iPSCs

To assess how X-dosage differentially primes mouse PSCs for rapid exit from pluripotency and to identify additional candidate regulators, we set out to globally define the open chromatin landscape of XX, XY and XO iPSCs. We employed assay for transposase-accessible chromatin (omniATAC-seq) to profile genome-wide chromatin accessibility with high resolution [34]. We generated ATAC-seq datasets from isogenic XX, XY and XO iPSC lines (and XX/XY ESCs), allowing to define the open chromatin regions and the enrichment for TF binding motifs associated with open chromatin landscapes (Fig 2, S4 Table). As expected, we observed open chromatin peaks at TSS-proximal and distal genomic regions, suggesting enrichment in cis-regulatory sequences (S2 Fig A). We could also use the mean read count ratio to autosome to infer the XX, XO and XY state of the cells (S2 Fig B). We then compared autosomal chromatin accessibility globally, and found that X-dosage impacts the chromatin accessibility landscape of iPSCs. Broadly, we observed a correlation between the number of active X-chromosomes and open chromatin landscapes (Fig 2A). We then assessed differential accessibility between XX and XY iPSCs, and between XO and XY iPSCs. We found that most open chromatin regions were shared between XX and XY iPSCs, suggesting that XX and XY iPSCs globally display similar open chromatin landscapes. However, thousands of chromatin regions showed increased accessibility in XX or in XY iPSCs, but not between XO and XY iPSCs (>2-fold, false discovery rate (FDR)<0.05, Fig 2B). These results further support the idea that X-dosage influences chromatin accessibility in iPSCs. We identified 2819 and 2363 autosomal chromatin regions that are more open in XX iPSCs or in XY iPSCs, respectively (Fig 2B, defined as "XX gain" and "XY gain" regions, S4 Table), which represent differences in chromatin accessibility driven by X-dosage. We also found a strong correlation between X-dosage and open chromatin landscapes in isogenic ESCs isolated from another genetic background (S2 Fig C/D, S5 Table). In summary, these results indicate that the chromatin landscape of XX and XY iPSCs is globally similar, but also contains differentially accessible chromatin at thousands of specific genomic regions, due to differences in X-dosage.

**Fig 2.**
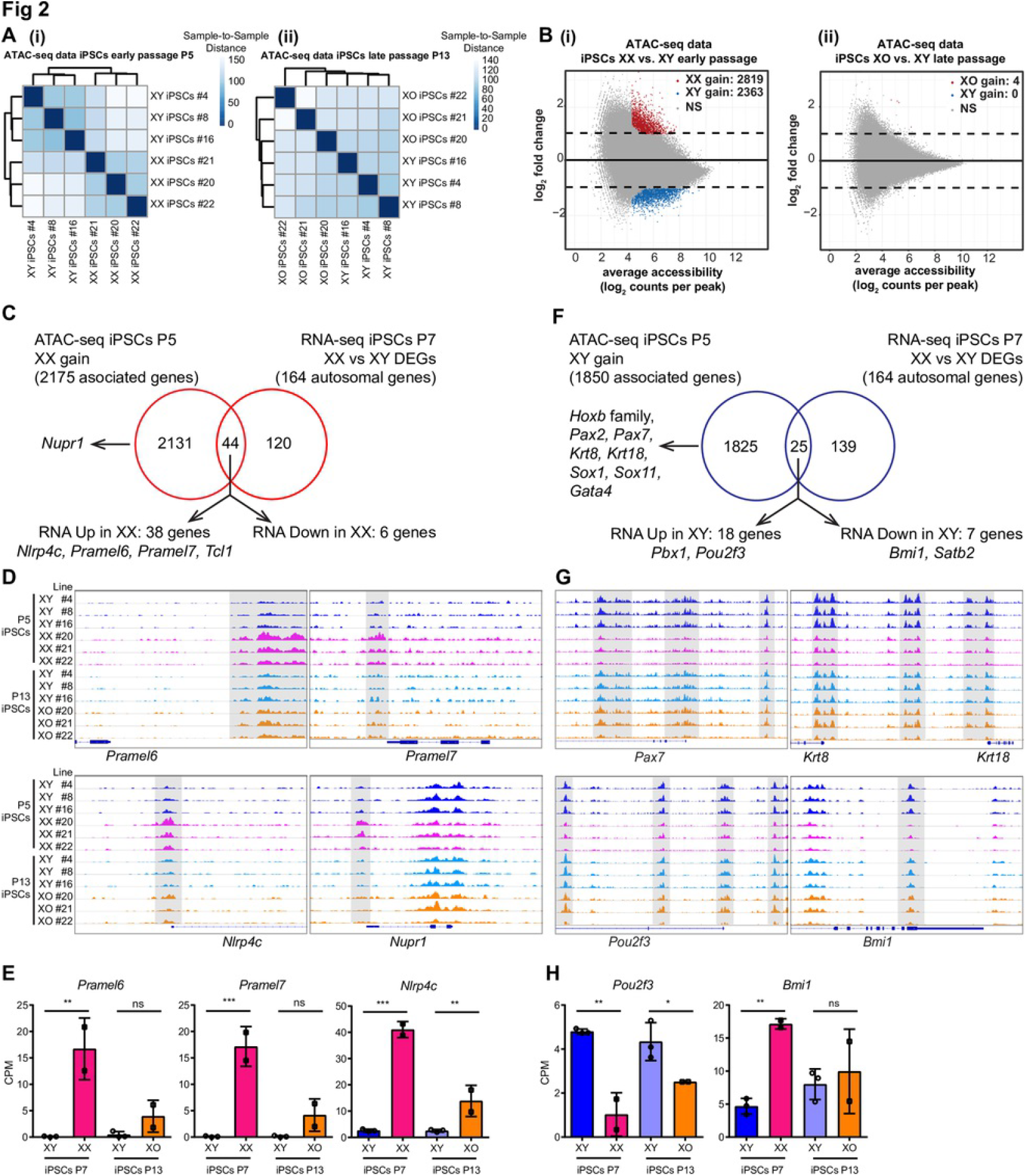
X-dosage modulates the chromatin regulatory landscape of mouse iPSCs. (A) ATAC-seq sample-to-sample distance heatmap showing the Euclidean distances (calculated from the rlog transformed counts, DESeq2) between iPSCs samples. (i) XX vs XY iPSCs, chromatin landscapes cluster by X-dosage (ii) XO vs XY iPSCs, the analysis cannot distinguish XO vs XY landscapes.
(B) Differential chromatin accessibility analysis between XX (or XO) and XY iPSCs. Log2 fold change (XX/XY or XO/XY) in reads per accessible region are plotted against the mean reads per ATAC-seq peak. Thousands of open chromatin regions are more open in XX iPSCs or in XY iPSCs (i), but not between XO and XY iPSCs (ii) (|log2fold|>=1, false discovery rate (FDR)<=0.05). These regions were defined as “XX gain” and “XY gain”, respectively.
(C) Venn diagrams showing the overlap between genes nearest to the “XX gain” regions and the DEGs between XX and XY iPSCs (DEGs= |log_2_fold|>=log_2_1.5, FDR<=0.05).
(D) Integrated Genome Viewer (IGV) track images of ATAC-seq signal for “XX gain” example regions in all iPSCs samples. Differentially open regions are shaded.
(E) Expression of *Pramel6*, *Pramel7* and *Nlrp4c* in XX, XY and XO iPSCs as assessed by RNA-seq. CPM value (count-per-million) were plotted for each gene. These genes are more expressed in XX iPSCs. Results are presented as averages (±SEM) of XY male, XX female iPSC lines in early passage (P7) and late passage (P13) XY and XO iPSCs, respectively. **p<0.01, ***p<0.001, P8 XX iPSCs vs P8 XY iPSCs. #p<0.05, ##p<0.01, P14 XO iPSCs vs P14 XY iPSCs. One-way ANOVA with Sidak’s multiple comparisons test.
(F) as in (C) for “XY gain” regions.
(G) As in (D) for “XY gain” regions”.
(H) as in (E) for *Pou2f3* and *Bmi1.*

We next assessed the correlation between differentially open chromatin and gene expression. Broadly, we observed a weak correlation between changes in chromatin accessibility and changes in gene expression (Fig 2C-F). Most differentially open chromatin regions did not associate with DEGs (2131/2175 genes for XX gain regions, 1825/1850 genes for XY gain regions, Fig 2C/F). Likewise, most DEGs did not associate with differentially accessible chromatin regions (120/164 and 139/164 DEGs were not associated with changes in chromatin accessibility in XX or in XY iPSCs, respectively, Fig 2 C/F). Nevertheless, a small fraction of differentially open chromatin regions associated with DEGs (Fig 2C/F, S6 Table). We identified 44 genes out of 164 autosomal DEGs that associated with chromatin regions more open in XX female iPSCs. Most of these genes (86%, 38/44) were transcriptionally upregulated in XX female iPSCs cells (Fig 2C). For example, there were chromatin regions more accessible in XX female iPSCs that associated with pluripotency-associated genes *Pramel6* and *Pramel7*, both of which were upregulated in XX female iPSCs, but not in XO female iPSCs (Fig 2C-E, S6 Table). Overexpression of *Pramel6* and *Pramel7* was found to oppose exit from pluripotency [35] and *Pramel7* was shown to mediate ground-state pluripotency [36]. We also observed increased accessibility in the vicinity of *Nlrp4c*, *Nupr1* and *Tcl1* in XX female iPSCs, but not XO female iPSCs (Fig 2C-E, S6 Table). These results indicate that the open chromatin landscape of iPSCs reflects specific cellular states, where XX-specific open chromatin could mediate stabilization of pluripotency in XX female iPSCs and ESCs.

Chromatin regions more accessible in XY male iPSCs associated with multiple genes involved in embryonic development and morphogenesis (several *Hoxb* genes, *Pax2*, *Pax7*, *Krt8, Krt18, Sox1, Sox11, Gata4*, S6 Table). 25 genes associated with chromatin regions more open in XY male iPSCs, 72% of which (18/25) were upregulated in XY male iPSCs (Fig 2F). Examples of upregulated genes include *Pou2f3* and *Pbx1* (Fig 2H). Within the 25 DEGs associated with XY gain chromatin regions in iPSCs, 7 genes were downregulated in XY iPSCs (*Bmi1*) (Fig 2F-H, S6 Table). In summary, these findings indicate that chromatin more open in XY or in XX iPSCs is associated with several lineage specification/ differentiation related and pluripotency-associated genes, respectively.

### Motif Analysis Reveals Potential Regulators of X-Dosage-Mediated Cell States

To identify TFs involved in modulating iPSCs as a result of differences in X-dosage, we searched for known TF motifs enriched in chromatin more open in XX female or in XY male iPSCs. Motif enrichment analysis of chromatin regions more open in XX female iPSCs revealed a strong enrichment for the binding motif of TFs such as RFX (10%), KLF5 (48.14%), ZIC3 (26.32%), and NANOG (26.92%) (Fig 3A). Regulatory factor X (RFX) proteins encode TFs expressed in many tissues including brain and testes [37]. KLF5 and NANOG have been functionally implicated in ESCs self-renewal (reviewed in [38]). ZIC3 is a pluripotency-associated factor required to maintain pluripotency [39]. Interestingly, *Zic3* is located on the X-chromosome, raising the possibility that ZIC3 dosage could drive X-linked driven stabilization of pluripotency in XX female iPSCs (see below). In summary, several top TF motifs enriched in chromatin with increased accessibility in XX female iPSCs belong to pluripotency-associated factors, suggesting that the identified pluripotency-associated TFs participate in stabilizing the pluripotency transcriptional regulatory network of XX female iPSCs.

**Fig 3.**
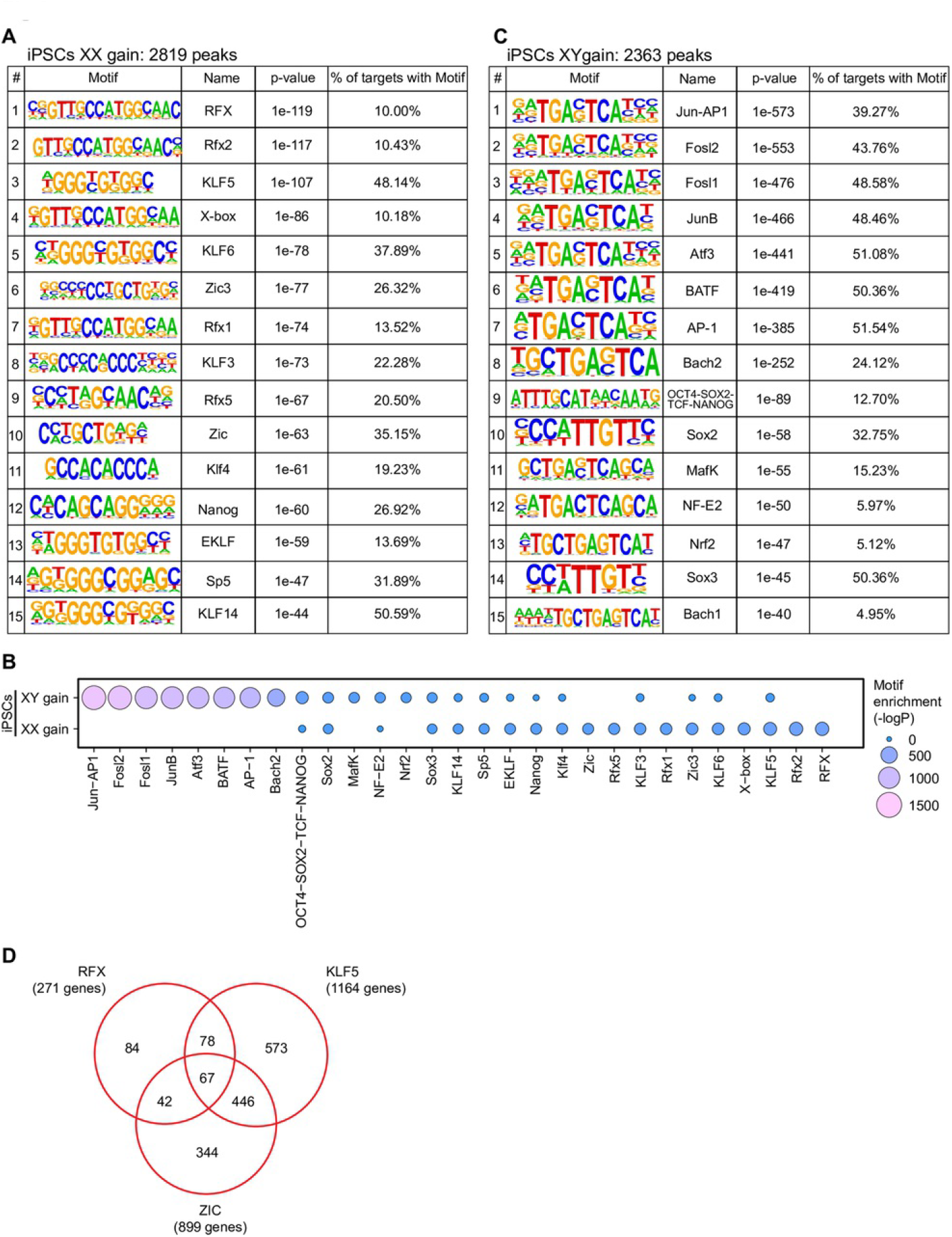
Identification of candidate regulators mediating the effects of X-dosage on open chromatin. (A-C) TF motifs enriched in chromatin regions more open in XX iPSCs or XY iPSCs.
(D) Venn diagram showing the overlap between genes associated the ATAC-seq regions more open in XX female iPSCs that contain a motif for KLF5, RFX or ZIC. The number of genes associated with all three motifs is indicated.

By contrast, pluripotency-associated TF motifs were less represented from the top motifs enriched in chromatin with increased accessibility in XY male iPSCs, with a few exceptions (Fig 3B/C). Instead, within chromatin more open in XY male iPSCs, motif enrichment analysis revealed binding motifs of multiple TFs of the AP-1 family such as JUN/AP-1 (39.27%), FOSL2 (43.78%) and ATF3 (51.08%) (Fig 3B/C). JUN/AP-1 is a transcriptional activator complex involved in regulating many processes [40,41] including cell growth and differentiation in response to a variety of stimuli including the MAPK pathway [42,43]. FOSL2 is a member of the AP-1 complex [40]. Collectively, these findings reveal that X-dosage modulates chromatin accessibility in iPSCs. As expected, we made similar observations in ESCs (S2 Fig C-I). In addition, open chromatin regions that are common between XX female and XY male iPSCs still showed enrichment of pluripotency-related TFs (S2 Fig I/J). We propose that the differential enrichment of TF binding sites in open chromatin regions modulated by X-dosage provides a molecular link between transcriptional regulators, stabilization of pluripotency in XX female PSCs and rapid exit from pluripotency in XY male PSCs.

In order to identify the putative target genes, we searched for genes associated with open chromatin regions enriched for specific motifs, then determined the target genes shared for open chromatin containing more than one motif. In chromatin more open in XX iPSCs, we found that 67 genes were associated with binding motifs for all three TF motifs RFX, KLF and ZIC (Fig 3D). Taken together, these analyses allowed the identification of TFs that regulate a large number of cis-regulatory regions, thereby improving our understanding on how X-dosage can drive two distinct pluripotent stem cell states.

### *Zic3* and *Tfe3* Dosage Do Not Explain X-Dosage Differences in Transcription and Pluripotency Exit

We sought to test if X-linked pluripotency-associated genes with enriched motifs identified in chromatin more open in XX female iPSCs stabilize pluripotency in XX female PSCs. Our motif discovery analysis identified the X-linked gene *Zic3* within the top motifs enriched in chromatin more open in XX female iPSCs and ESCs (Fig 3A, S2 Fig F). Western blot analysis showed that XX iPSCs and ESCs express higher ZIC3 protein than XY male iPSCs and ESCs (Fig 4A). In addition, the increased *Zic3* transcript levels of XX female iPSCs were restored to XY male levels in XO female iPSCs (S3 Fig A). Moreover, *Zic3* was reported to prevent endodermal lineage specification and to act as a transcriptional activator of *Nanog* expression [39,44], further suggesting that it could have a role in stabilizing pluripotency in XX female ESCs. To test the hypothesis that increased *Zic3* dosage stabilizes pluripotency in XX female ESCs, we overexpressed *Zic3* in XY male iPSCs and asked if it induced XX-like features (S3 Fig B). We achieved 3 fold overexpression of ZIC3 protein tagged in N- or C-terminal with HA (S3 Fig C). We then subjected the cells to LIF withdrawal. We found that overexpression of *Zic3* with a N-terminal HA-tag, but not with the C-terminal HA-tag, delayed pluripotency exit during LIF withdrawal differentiation (S3 Fig B-E). These results suggested that increased *Zic3* dosage might be responsible for the pluripotency exit delay of XX female PSCs. Using an independent approach, we generated *Zic3* heterozygous deletions using CRISPR-Cas9 genome editing in XX female ESCs to reduce *Zic3* dosage, which is a more stringent method to test whether *Zic3* dosage stabilizes pluripotency in XX female ESCs (Fig 4B-D, S3 Fig F/G). Two independent *Zic3+/-* XX female ESC clones maintained XX-like expression of *Prdm14*, *Nanog* and *Tcl1* and also maintained female-like delayed exit from pluripotency (Fig 4E). We performed similar experiments for another additional X-linked gene involved in pluripotency, *Tfe3*, and obtained similar results as for *Zic3* (Fig 4F-H, S3 Fig H-J). These results support the idea that the dosage of *Zic3* and *Tfe3* do not explain the differences in pluripotency gene expression and exit from pluripotency between XX female and XY male ESCs.

**Fig 4.**
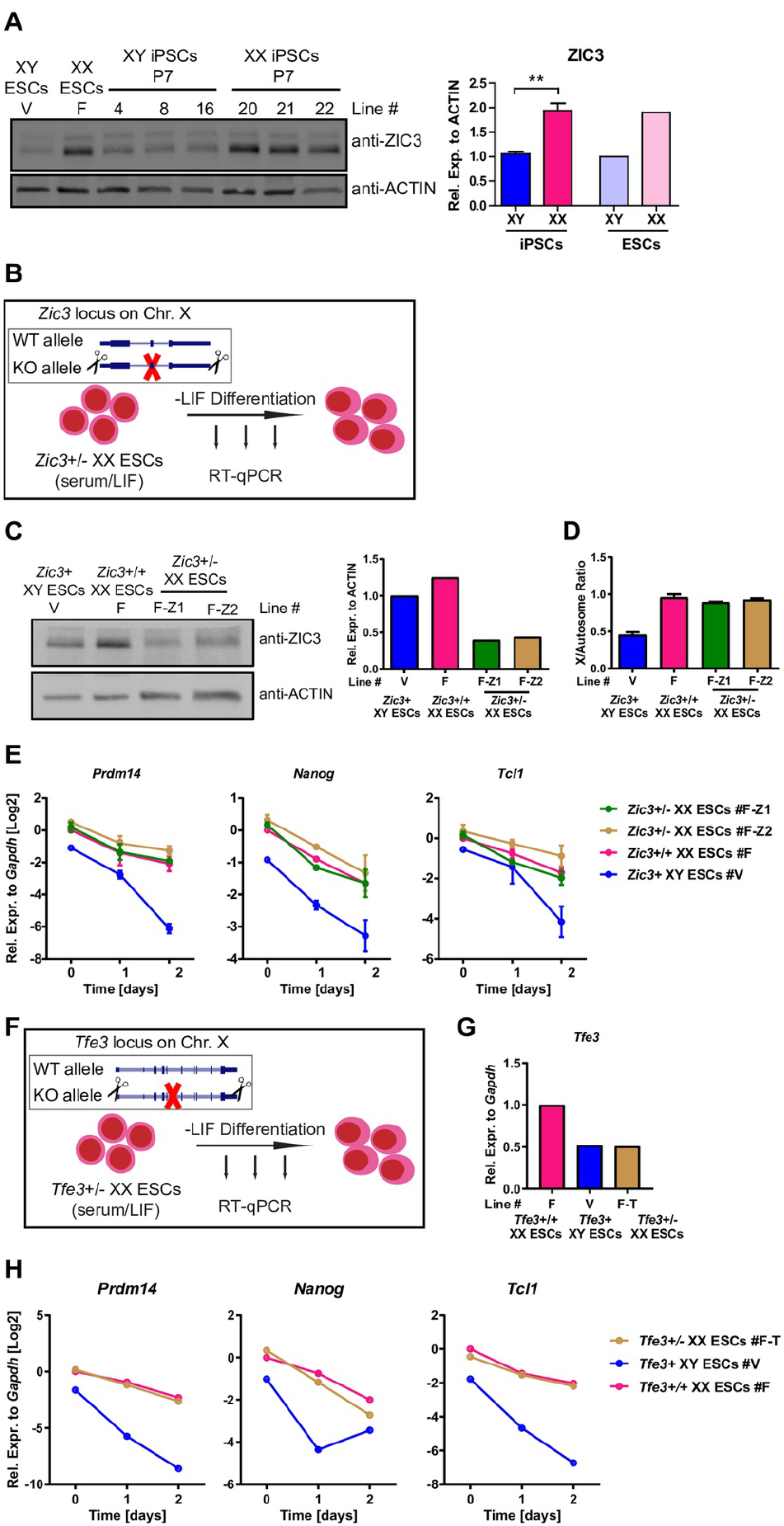
Effects of Zic3/Tfe3 heterozygous deletion on pluripotency exit. (A) Western blot analysis for ZIC3 in iPSCs and ESCs grown in S/L. Right: quantification using ACTIN as loading control. ZIC3 protein values are represented as averages (±SEM) of XY and XX iPSC lines (three lines each), and XY and XX ESC lines (one line each). **p<0.01, XY iPSCs vs XX iPSCs, unpaired two-tailed t-test.
(B) Scheme of heterozygous *Zic3* deletion strategy in XX female ESCs followed by LIF withdrawal.
(C) Western blot analysis for ZIC3 in two independent *Zic3+/-* ESC lines, *Zic3+/+* ESCs and XY ESCs all grown in S/L (n=1).
(D) qPCR analysis for X-chromosome DNA copy number. X copy number are presented as the average ratio of gDNA quantities for four X-linked genes (*Tfe3, Bcor, Pdha1, and Mid1*) to gDNA quantities for autosomal gene *Gapdh.* Results are presented as averages (±SEM) of the same cell lines in two independent qPCR experiments (n=2).
(E) RT-qPCR analysis for pluripotency-associated gene expression during LIF withdrawal in the two independent *Zic3+/-*XX ESC lines, the *Zic3+/+* XX parental ESC line and a XY ESC line. Results are presented as averages (±SEM) of the same lines in two independent differentiation experiments (n=2).
(F) Scheme of heterozygous *Tfe3* deletion strategy in XX female ESCs followed by LIF withdrawal.
(G) RT-qPCR analysis for *Tfe3* expression in the *Tfe3+/-* XX ESC line, the *Tfe3+/+* XX parental ESC line and a XY ESC line (n=1).
(H) RT-qPCR analysis for pluripotency-associated gene expression during LIF withdrawal in the *Tfe3+/-* XX ESC lines, the *Tfe3+/+* XX parental ESC line and a XY ESC line. Results are presented as averages (±SEM) of biological duplicates.

### *Dkc1*, *Otud6a*, *Fhl1*, *Zfp185* and *Scml2* Dosage Do Not Explain X-Dosage-Specific Differences in Pluripotency Exit

We sought to find the X-linked regulators that drive stabilization of pluripotency in XX female PSCs. We analyzed RNA-seq and published proteomics data of XX female and XY male ESCs [17]. We selected X-linked candidate factors with 1) increased expression in XX ESCs over XY or XO ESCs and ranked by expression ratio, 2) evidence that the genes are subject to X-chromosome inactivation (S7 Table), and 3) literature consistent with a possible role in stabilizing pluripotency. The selected candidate genes were *Dkc1, Otud6a, Fhl1, Zfp185* and *Scml2*. We overexpressed their cDNAs in XY male iPSCs (S4 Fig). To test the effect of overexpression on pluripotency exit, we induced differentiation by LIF withdrawal and measured pluripotency gene expression at 24h and 48h. We found that overexpression of *Dkc1, Otud6a, Fhl1, Zfp185* or *Scml2* was not sufficient to induce a delay in pluripotency exit (S4 Fig). Collectively, these findings do not support a significant role for these X-linked pluripotency-associated genes in stabilizing pluripotency in XX female ESCs.

### Heterozygous *Dusp9* Deletion in XX Female ESCs Induces Male-Like DNA Methylation Yet Maintains A Female-Like Transcriptome, Open Chromatin Landscape, Growth and Delayed Pluripotency Exit

In an effort to understand the mechanisms by which X dosage affects pluripotent stem cell properties, we generated *Dusp9* heterozygous deletions in XX female ESCs using CRISPR-Cas9 genome editing, resulting in two independent *Dusp9+/-* XX female ESC clones (Fig 5A/B, S5 Fig A/B). To ensure the maintenance of two active X-chromosomes in *Dusp9* +/- ESCs, we used polymorphic *Musculus/Castaneus* (Mus/Cas) ESCs, known to be less susceptible to X-chromosome loss [17,45]. We confirmed that *Dusp9*+/- ESCs maintained two active X-chromosomes (S5 Fig C/D). *Dusp9+/-* XX female ESCs showed male-like global DNA methylation (S5 Fig E), corroborating recent findings [17]. To determine if *Dusp9*+/- XX female ESCs with male-like DNA methylation acquire male-like transcription, we analyzed the expression of stem cell maintenance related genes using RNA-seq in *Dusp9*+/- XX female ESCs, *Dusp9*+/+ XX female ESCs and *Dusp9*+ XY male ESCs, all sharing a Mus/Cas background to exclude potential strain-specific effects. Principal component analysis (PCA) placed *Dusp9+*/- XX female ESCs away from both *Dusp9+/+* XX female ESCs and *Dusp9*+ XY male ESCs, indicating that *Dusp9*+/- XX female ESCs do not adopt a male-like transcriptional state (Fig 5C). We corroborated this finding using unsupervised clustering of stem cell maintenance related gene expression, where *Dusp9*+/- XX female ESCs clustered together with *Dusp9+/+* XX female ESCs, and away from *Dusp9*+ XY male ESCs (Fig 5D). Unsupervised clustering analysis also showed the activation of most MAPK target genes in *Dusp9*+/- XX female ESCs, in agreement with the function of *Dusp9* as a MAPK inhibitor (Fig 5E) [46]. Furthermore, we found more DEGs between *Dusp9*+/- XX female ESCs and XY male ESCs, and less DEGs between *Dusp9*+/- XX female ESCs and *Dusp9*+/+ XX female ESCs, with little overlap between the two sets of DEGs (Fig 5F/G). The only exception was the *Pramel7* gene, the expression of which was reduced to XY male levels in *Dusp9*+/- XX female ESCs, indicating that transcription of *Pramel7* is influenced by *Dusp9* dosage (S8 Table). Overall these results indicate that, male-like DNA methylation can be induced in the absence of male-like transcription in *Dusp9+/-* XX female ESCs. These experiments raise the possibility that distinct molecular features modulated by X-dosage in ESCs might be controlled by different regulators.

**Fig 5.**
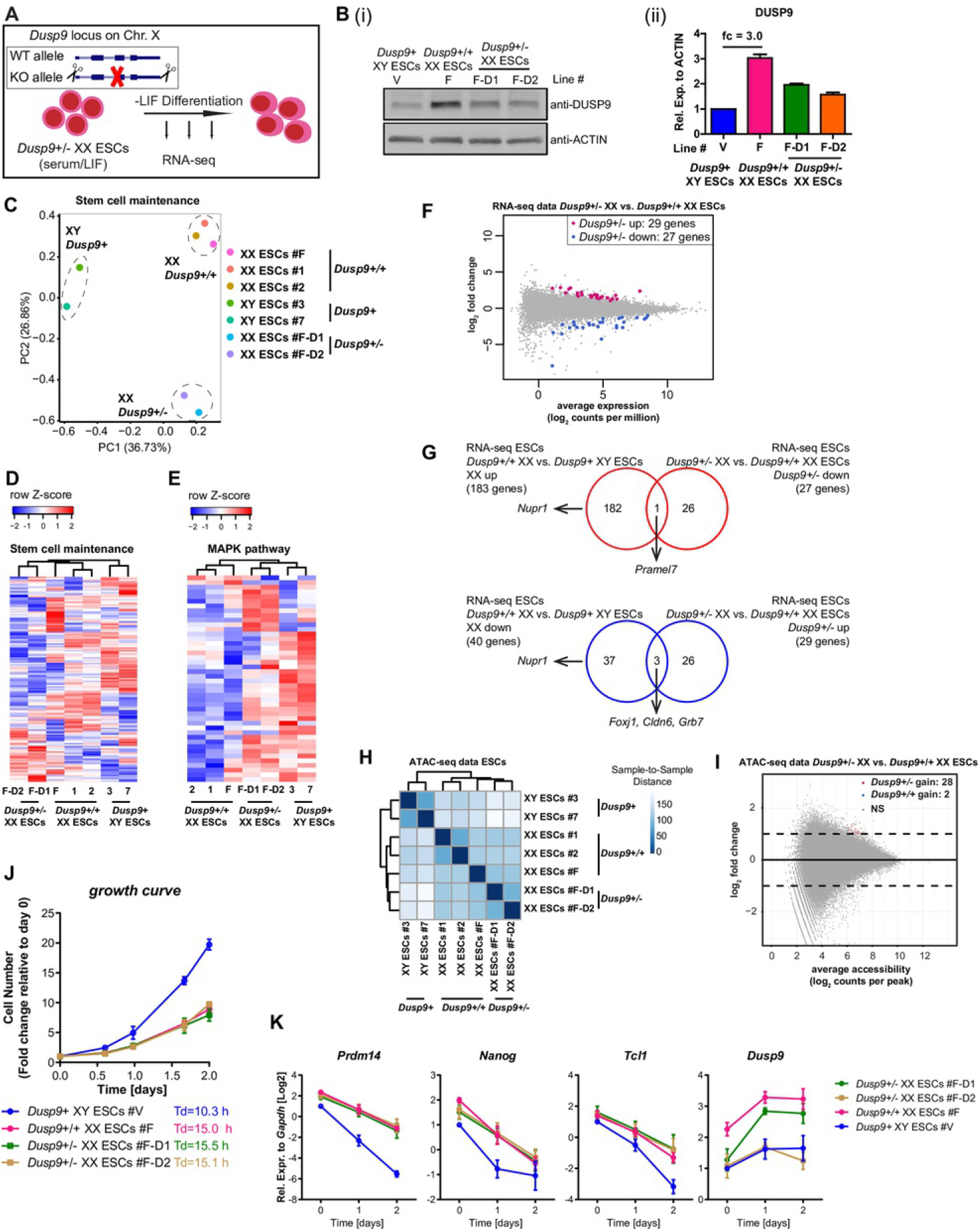
The open chromatin landscape, growth and pluripotency exit delay of XX ESCs are maintained in the presence of global DNA methylation. (A) Scheme of *Dusp9* heterozygous deletion in XX female ESCs followed by LIF withdrawal differentiation.
(B) (i) Western blot analysis for DUSP9 in *Dusp9+/-* ESCs, *Dusp9+/+* ESCs and XY ESCs grown in S/L. (ii) Quantification of DUSP9 levels using ACTIN as a loading control. DUSP9 expression values are represented as averages (±SEM) of the same lines in two independent western blot experiments.
(C) PCA of RNA-seq data for *Dusp9+/-* ESCs, *Dusp9+/+* ESCs and XY ESCs grown in S/L conditions. Stem cell maintenance genes (GO:0019827) were used in this analysis.
(D) Unsupervised hierarchical clustering of stem cell maintenance genes (GO:0019827) for *Dusp9* +/- ESCs, *Dusp9* +/+ ESCs and XY ESCs in the undifferentiated state.
(E) Unsupervised hierarchical clustering of MAPK pathway related genes (defined in [15]) for *Dusp9+/-* ESCs, *Dusp9* +/+ ESCs and XY ESCs in the undifferentiated state.
(F) DEG analysis, identifying clear differences between *Dusp9+/+* XX ESCs and XY ESCs, but much less between *Dusp9* +/- XX EScs and *Dusp9+/+* XX ESCs.
(G) Overlap between the DEGs of *Dusp9+/+* vs XY ESCs and *Dusp9* +/- vs *Dusp9* +/+ ESCs for upregulated genes (up) and downregulated genes (down), (|log_2_fold|>=log_2_1.5, FDR<=0.05). *Dusp9* heterozygous deletion maintains a female-like transcriptome.
(H) ATAC-seq sample-to-sample distance heatmap showing the Euclidean distances (calculated from the rlog transformed counts, DESeq2) between open chromatin landscapes in *Dusp9* +/+, *Dusp9* +/- and XY ESCs. *Dusp9* +/- ESCs maintain a *Dusp9*+/+-like open chromatin landscape.
(I) Differential chromatin accessibility analysis between *Dusp9* heterozygous mutant and wild type XX ESCs. Log2 fold change (mutant/wild type) in reads per accessible region are plotted against the mean reads per ATAC-seq peak. *Dusp9* heterzygous mutants maintain a female-like open chromatin landscape. (|log2fold|>=1, false discovery rate (FDR)<=0.05).
(J) Growth curves and doubling times of *Dusp9+/-* ESCs, *Dusp9*+/+ ESCs and XY ESCs in S/L condition. Cells were counted at the indicated time points and presented as fold changes relative to averages (±SEM) at 0h. Data represent a representative experiment (from at least four independent experiments).
(K) RT-qPCR for *Prdm14*, *Nanog, Tcl1 and Dusp9* expression before and after LIF withdrawal. Results are presented as averages (±SEM) of three independent experiments.

Next, we sought to determine if chromatin accessibility is affected by reduced DUSP9 dosage in XX female ESCs. ATAC-seq revealed that *Dusp9*+/- XX female ESCs maintain a female-like open chromatin landscape (Fig 5H). We only observed very few differences in chromatin accessibility between *Dusp9*+/- XX female ESCs and *Dusp9*+/+ XX female ESCs (Fig 5I), while the same analysis identified thousands of regions differentially accessible in XX and XY ESCs (S2 Fig D). This finding indicate that the effects of X-dosage on chromatin accessibility can be by large dissociated from global DNA methylation levels.

We then measured the growth of *Dusp9*+/- XX female ESCs, and found that the cells grew as slow as their parental *Dusp9*+/+ XX female ESCs, both of which grew slower than XY male ESCs (Fig 5J, Fig 1K). Therefore, reducing *Dusp9* dosage and inducing global DNA methylation in XX female ESCs is not sufficient to induce male-like cellular growth. We propose that X-dosage-specific growth and global DNA methylation are regulated by different pathways in mouse PSCs.

To study the effects of *Dusp9* heterozygous deletion in XX female ESCs on pluripotency exit, we subjected *Dusp9*+/- XX female ESCs, *Dusp9*+/+ XX female ESCs and XY male ESCs to LIF withdrawal differentiation for 48h followed by RT-qPCR analysis. The delay in pluripotency exit as judged by *Prdm14, Nanog* and *Tcl1* expression was maintained in *Dusp9+/-* XX female cells relative to *Dusp9*+/+ XX female cells (Fig 5K). In further support of the finding that *Dusp9*+/- XX female ESCs maintain a delay in pluripotency exit, RNA-seq analysis showed that multiple pluripotency-associated genes behaved similarly in *Dusp9*+/- XX female and *Dusp9*+/+ XX female cells undergoing differentiation (S5 Fig F). Therefore, mechanistically, reducing *Dusp9* dosage is compatible with female-like pluripotency exit. We conclude that reducing the dosage of *Dusp9* in XX female ESCs is not sufficient to induce a male-like transcriptome or accelerate pluripotency exit to a male-like state, despite changes in the expression level of multiple genes in the MAPK signaling pathway and despite male-like DNA methylation. In addition, *Dusp9* overexpression in XY male ESCs did not induce a female-like delay in differentiation (S5 Fig G-L) despite inducing female-like global DNA hypomethylation [17].

Altogether, these results indicate that most changes in open chromatin, growth and pluripotency exit as a result of differences in X-dosage are regulated independently of global DNA methylation in XX ESCs. Hence, mechanistically, heterozygous *Dusp9* deletion molecularly uncouples global DNA methylation from the open chromatin landscape, growth, and the pluripotency exit delay of XX female ESCs. Importantly, this points towards the existence of other pathways and X-linked genes involved in mediating the effects of X-dosage in PSCs.

### Large-Fragment Heterozygous Deletions

Two models emerged to mechanistically explain delayed pluripotency exit in XX PSCs. In the first model, a single X-linked gene is responsible for delayed pluripotency exit. In the second model, multiple X-linked genes act together to delay pluripotency exit. To test these models, we generated a series of large fragment (LF) heterozygous deletions of the X chromosome in XX ESCs, which were confirmed by genotyping and Sanger sequencing (S6 Fig), and then induced to differentiate. There was a partial rescue of the pluripotency exit delay of XX ESCs in multiple, but not all, LF deletions (Fig 6, S1 Table). The partial rescue was gene-specific and fragment-specific. These results favor a mechanism corresponding to model 2, where multiple X-linked genes participate in delayed pluripotency exit in XX PSCs.

**Fig 6.**
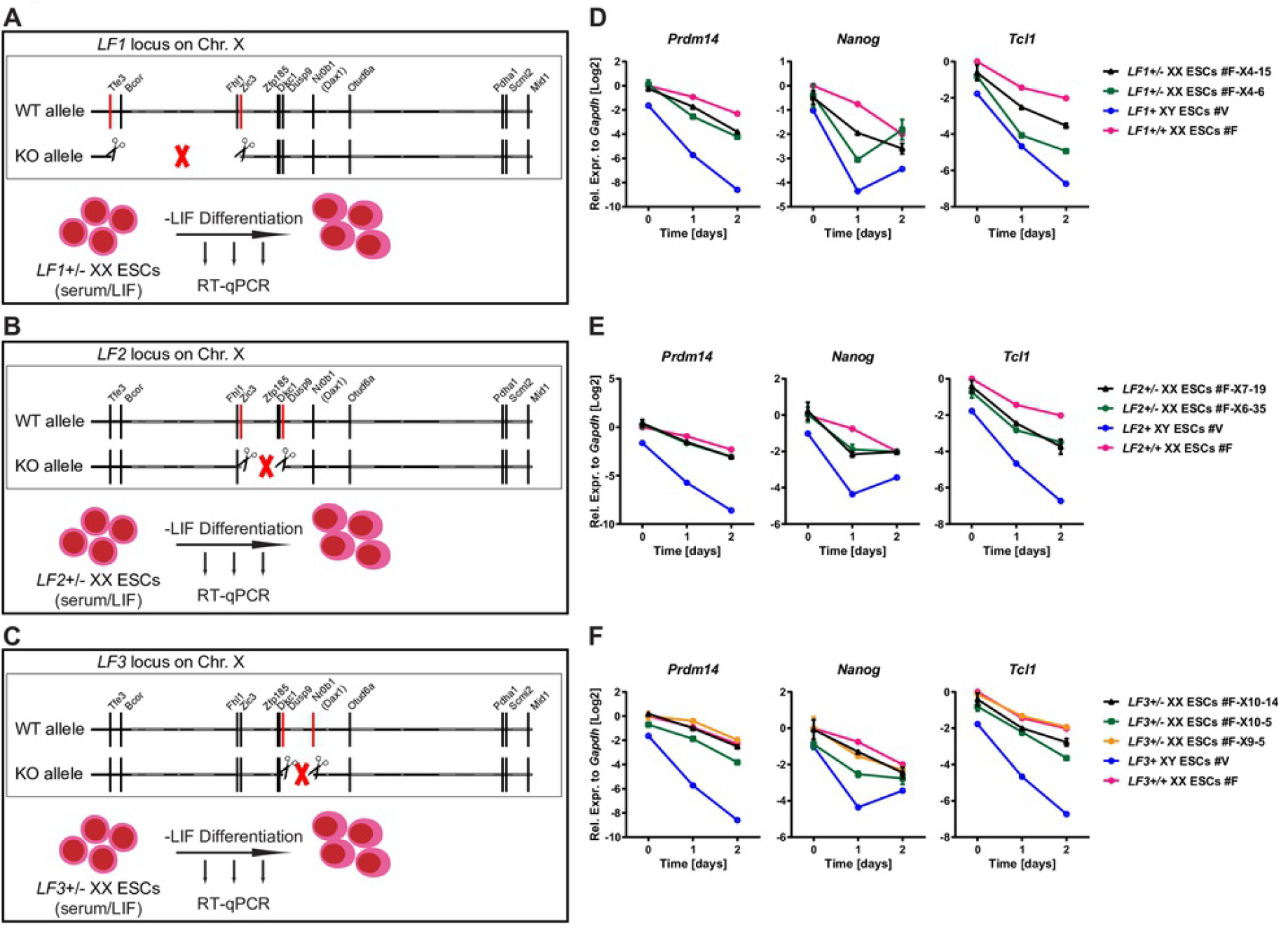
Multiple X-linked genes modulate the pluripotency exit delay of XX PSCs. (A-C) Scheme of large fragment (LF) heterozygous deletions in XX female ESCs followed by LIF withdrawal differentiation. (A) LF1, (B) LF2, (C) LF3.
(D-F) RT-qPCR for *Prdm14*, *Nanog and Tcl1* expression before and after LIF withdrawal in two-three independent LF+/-XX ESC lines, the parental XX ESC line and a XY male ESC line. Results are presented as averages (±SEM) of biological duplicates. (D) LF1, (E) LF2, (F) LF3.

## DISCUSSION

Induction of naive pluripotency during reprogramming to iPSCs and during *in vivo* development in inner cell mass (ICM) leads to X-chromosome reactivation in murine female cells [6,7,30,47]. The consequences of X-dosage imbalance between female (XX) and male (XY) cells on mouse iPSCs remained unclear until now. In addition, the regulatory mechanisms at the basis of distinct X-dosage-specific features in mouse PSCs were incompletely understood. In this study, we addressed these questions by analyzing the transcriptome, growth properties, chromatin accessibility landscape and pluripotency exit of isogenic female and male iPSCs. We identified X-dosage as a factor influencing the molecular and cellular properties of iPSCs. By employing epigenomic analyses we found that X-dosage modulates open chromatin in iPSCs and ESCs. Furthermore, using genome editing we found that modulation of the transcriptome, open chromatin, cell growth and pluripotency exit by X-dosage is by large independent of global DNA methylation. We provide evidence favoring a model where multiple X-linked genes delay pluripotency exit. Our results change the concept of how X-dosage influences pluripotency.

### Impact of X-Dosage on the Transcriptome and Pluripotency Exit

One outcome of our study is that the number of active X-chromosomes correlates with differences in the transcriptome and in pluripotency exit in iPSCs (Fig 1), in addition to differences in DNA methylation [22,23]. Reprogramming somatic cells to iPSCs is an important system to study erasure of epigenetic memory and pluripotency. Sex does not appear to influence the efficiency of iPSCs generation, since we previously showed that female and male cells reprogram with similar efficiencies in this system [27]. However, we have now established that the presence of two active X-chromosomes, as a result of reprogramming to pluripotency in female cells, is associated with the slower growth of XX iPSCs, an altered transcriptome including increased pluripotency-associated gene expression, and delayed pluripotency exit. These differences are caused by changes in X-dosage, since XO iPSCs revert to an XY male iPSC phenotype. While this is the first study, to our knowledge, to report the influence of X-dosage on the growth of mouse PSCs, and on the transcriptome and pluripotency exit of iPSCs, our results are consistent with studies in mouse ESCs [10,13,15,18], in human ESCs [11,12,48] and in postimplantation mammalian embryos [10,32,49]. However, X-dosage is currently largely ignored in most reprogramming studies, in which neglecting sex-specific differences could have a negative impact on results interpretation. The notion that X-dosage influences the molecular and cellular properties of iPSCs [10,15,22] is further supported by the loss of sex-specific differences concomitant with loss of one X-chromosome in female iPSCs, in agreement with previous observations in ESCs [13,15,17,18] and in iPSCs [23]. The important point is that studies of reprogramming to iPSCs should consider the number of active X-chromosomes as a modulator of the transcriptome, and cells of different sex should be studied separately, but also often both considered.

The influence of X-dosage on the heterogeneity of ESCs also remained unclear. Our analysis of single cell RNA-seq data for XX and XY ESCs in S/L and 2i/L [24] revealed that both XX and XY ESCs reside in a metastable state, with *Nanog* high and *Nanog* low cells (S7 Fig). More XX ESCs express *Tcl1* than XY ESCs, in agreement with increased pluripotency-associated gene expression in XX ESCs [15]. Differences between the transcriptome of XX and XY ESCs persist in 2i/L, despite more homogenous pluripotency-associated gene expression (S7 Fig). Together, this analysis reveals that XX PSCs in S/L maintain a metastable state with a bias toward increased expression of specific pluripotency-associated genes, whereas transcriptomic differences between XX and XY PSCs persist in 2i/L.

### Impact of X-Dosage on Cell Growth and Cell-Cycle Control

Here we show for the first time that the presence of two active X-chromosomes in iPSCs and ESCs is associated with delayed cellular growth. One possible interpretation is that the delayed growth of female post-implantation mammalian embryos [32,49] is recapitulated in mouse PSCs. Since the growth differences of XX and XY/XO PSCs are maintained after dual GSK3B and ERK inhibition, additional pathways are likely involved. One hypothesis is that there could be a competitive growth advantage of cells that have undergone X-chromosome inactivation in the post-implantation mammalian embryo to select against remaining cells that may fail to undergo X-chromosome inactivation, and maintain two active X-chromosomes. Our *in vitro* experiment suggests that X-chromosome inactivation could indeed provide a small growth advantage. However, this hypothesis remains to be tested *in vivo*.

### Differences in Chromatin Accessibility Underlie X-Dosage Dependent Pluripotency States

To better understand what drives the features of the pluripotent regulatory network that are modulated by X-dosage in PSCs, we explored the open chromatin landscapes of female and male iPSCs and ESCs. While both female and male iPSCs/ESCs possessed globally similar open chromatin landscapes, thousands of chromatin regions were differentially accessible in XX and XY PSCs (Fig 2). These differentially accessible regions may underlie differences in the transcriptional regulatory network and functional properties of XX and XY PSCs. Decoding differentially accessible chromatin regions, we identified pluripotency-associated genes *Pramel6* and *Pramel7* with increased accessibility in XX iPSCs. *Pramel7* has been associated with naive pluripotency [36], and overexpression of *Pramel6* and *Pramel7* both compromise pluripotency exit [35]. We went further by identifying a catalogue of cis-regulatory regions including promoters that are modulated by X-dosage in iPSCs and ESCs. These observations indicate that X-dosage can modulate chromatin accessibility in mouse PSCs.

### Transcriptional Regulators of X-Dosage-Specific Pluripotent Stem Cell States

Decoding differentially accessible chromatin allowed us to distinguish distinct sets of enriched TF binding motifs in XX and XY ESCs. Specifically, motifs for KLF5, ZIC3 and NANOG were enriched in chromatin more open in XX iPSCs, all of which have been implicated in pluripotency [38,39,50–52]. These results suggest that the stabilization of pluripotency in XX female ESCs may be mediated by these core master regulators. In particular, *Zic3* is a known pluripotency factor [39], encoded on the X-chromosome, and is not dosage compensated in XX PSCs. However, *Zic3* heterozygous deletion had no effect on stabilization of pluripotency. Although no TF ChIP-seq data are available for XX iPSCs to date, our *in silico* analyses identified a high confidence set of direct putative KLF5, ZIC3, and NANOG targets in XX iPSCs, including known pluripotency-associated genes. Moreover, the specific putative regulatory region associated with *Pramel6* which becomes more accessible in XX iPSCs overlaps with ChIP-seq binding sites of OCT4, SOX2 and NANOG in XY ESCs (not shown), further suggesting that increased binding of master pluripotency regulators takes place at these more accessible regions in XX iPSCs. Our results raise the possibility that pluripotency is stabilized in XX iPSCs by binding of core pluripotency factors to a subset of regulatory elements whose accessibility is influenced by X-dosage.

In contrast to the XX state, chromatin more open in XY iPSCs identified AP-1 TFs as candidate regulators, which have not been implicated in X-dosage-specific regulation of pluripotency before. JUN/AP-1 control many cellular processes including proliferation, apoptosis and differentiation in response to a variety of stimuli including the MAPK pathway (reviewed in [40]). The role of AP-1 TFs in the context of X-dosage in iPSCs warrants future studies.

### Impact of *Dusp9* Dosage on the Molecular and Cellular Properties of ESCs

A previous study showed that *Dusp9* modulates DNA hypomethylation and the proteome in XX female mouse ESCs [17]. However, the effects of reducing *Dusp9* dosage in XX ESCs on growth, transcription and pluripotency exit were unknown. An important outcome of our analyses is that *Dusp9* heterozygous XX female ESCs maintain a female-like open chromatin landscape, growth and delayed pluripotency exit concomitant with male-like global DNA methylation levels (Fig 5). These results suggest that chromatin accessibility, growth and delayed pluripotency exit can be regulated independently of global DNA methylation levels in mouse PSCs. This result was unexpected for two reasons. First, *Dusp9* overexpression in ESCs was reported to induce a female-like proteome, including activation of naive pluripotency marker PRDM14 [17]. Second, reducing the expression of DNMTs in male ESCs is associated, at least in part, with delayed pluripotency exit [15]. However, it was reported that the ICM of female and male embryos show comparable DNA methylation [17], despite delayed female development, suggesting that DNA hypomethylation and stabilization of pluripotency can be uncoupled both *in vivo* and *in vitro*. Our results therefore suggest that global DNA methylation levels are regulated, at least in part, by distinct X-linked genes, different from those regulating the open chromatin landscape and stabilization of pluripotency in PSCs (*Dusp9* for DNA methylation levels, other gene(s) for chromatin accessibility and delayed pluripotency exit and growth). Choi et al. reported that *Dusp9* overexpression in male ESCs increases the expression of PRDM14, ROR2 and TFCP2L1 [17]. In our study, 3.5-3.7 fold overexpression of DUSP9 protein in male ESCs, achieving comparable DUSP9 protein level as in XX ESCs, did not lead to an increase in *Prdm14* transcript level. This may be explained by differences in the level or method of *Dusp9* overexpression (inducible system vs piggybac), or the assay used to judge expression of pluripotency markers (mass spectrometry vs RT-qPCR). Interestingly, *Pramel7* overexpression in ESCs was shown to induce DNA hypomethylation through the degradation of DNA methylation maintenance factor UHRF1 [36]. At the same time, we found that reducing *Dusp9* dosage in XX female ESCs reduces *Pramel7* expression to male levels. The results suggest that *Pramel7* may be downstream of *Dusp9*, and may participate in the control of DNA methylation by X-dosage.

### Multiple X-linked genes modulate pluripotency exit delay in XX PSCs

Our results do not support a model where a single X-linked gene stabilizes pluripotency in XX PSCs. First, single heterozygous deletions of *Zic3, Tfe3, Dkc1, Otud6a, Fhl1, Zfp185* and *Scml2* had little effect. Second, distinct large fragment heterozygous deletions suggested that multiple X-linked genes participate in delaying pluripotency exit in XX PSCs. Therefore, identifying additional X-linked regulators that mediate the effects X-dosage in pluripotent stem cells requires future investigations [10]. An interesting additional candidate is the recently-identified X-linked transient octamer binding factor 1 (TOBF1) [53], since it was shown to sustain pluripotency. It is also possible that other regulators of the Erk pathway are involved. A previous study in human ESCs reported that human primed PSCs with eroded X-chromosome inactivation and increased expression of the MAPK/ERK downstream effector ELK-1 have decreased expression of TRA-1-60, a marker of the differentiated state [12]. However, human primed PSCs studies are likely not compatible with mouse naive PSCs studies because cells reside in distinct pluripotent states.

To conclude, we revealed that global DNA methylation can be uncoupled from delayed pluripotency exit in XX mouse ESCs. Furthermore, our study shows for the first time that X-dosage-specific differences in cell growth, open chromatin landscape, transcription, and pluripotency exit in iPSCs correlate with the number of active X-chromosomes. We also reveal a mechanism by which multiple genomic regions on the X chromosome are responsible for delaying pluripotency exit. Using information from the genome, the epigenome and the transcriptome we gained insights into modulation of the open chromatin landscape and the transcriptional regulatory network of iPSCs by X-dosage. Furthermore, better understanding how X-dosage modulates pluripotency will have important implications for disease modeling and regenerative medicine.

## EXPERIMENTAL PROCEDURES

### Mice and reprogramming

MEFs were isolated from individual E14.5 mouse embryos obtained from a cross between wild type (WT) C57BL/6 and homozygous Rosa26:M2rtTA, TetO-OSKM mice [25]. Individual embryos were genotyped for sex using *Ube1* as previously described (See S9 Table for primer sequence) [27] using homemade Taq DNA Polymerase and grown in MEF medium [DMEM (Gibco, 41966-052) supplemented with 10% (v/v) fetal bovine serum (FBS, Gibco, 10270-106), 1% (v/v) penicillin/streptomycin (P/S, Gibco, 15140-122), 1% (v/v) GlutaMAX (Gibco, 35050-061), 1% (v/v) non-essential amino acids (NEAA, Gibco, 11140-050), and 0.8% (v/v) beta-mercaptoethanol (Sigma, M7522)]. Reprogramming was induced by doxycycline (final 2 μg/ml) in mouse ESC medium [KnockOut DMEM (Gibco, 10829-018) supplemented with 15% FBS, 1% (v/v) P/S, 1% (v/v) GlutaMAX, 1% (v/v) NEAA, 0.8% (v/v) beta-mercaptoethanol, and mouse LIF] in the presence of ascorbic acid (final 50 μg/ml). Individual colonies were picked at day 16 onto irradiated male feeders in ESC medium without doxycycline or ascorbic acid and expanded for three passages, eventually obtaining 10 female iPSC lines (lines 1, 4, 5, 6, 8, 12, 13, 14, 16, 17, 18) and 11 male iPSC lines (lines 19, 20, 21, 22, 23, 24, 25, 26, 27, 28, 29) at passage (P) 4 (S1 Fig B). iPSC lines 1, 4, 8, 12, 14, 16, 20, 21, 22, 24, 26 and 28 were also used in another study [23] (S1 Table). Mus/Cas ESCs were isolated from E3.5 embryos resulting from a cross between Cast/Eij males and C57B6/J females, as described [54]. All animal work carried out in this study is covered by a project license approved by the KU Leuven Animal Ethics Committee.

### Cell lines and culture

XY and XX Mus/Cas ESCs were newly derived in our lab and also obtained from the Deng laboratory [24]. XY ESCs (V6.5) and XX ESCs (F1-2-1) were obtained from the Plath laboratory. GFP-labelled (Oct4-GiP) XY ESCs were previously described [55]. ESCs and iPSCs (male iPSC line 4, 8, 16; female iPSC line 20, 21, 22) were expanded on top of male WT feeders in mouse ESC medium (S/L condition), eventually early passage cells (iPSCs: P6-P8) and late passage cells (iPSCs: P13-P14) were used for further experiments. ESCs and iPSCs (male iPSC lines 1, 4, 8, 12, 16; female iPSC lines 19, 20, 21, 22, 23, 26) were adapted to 2i/LIF, where cells grown on feeders in S/L condition (iPSCs: P4) were switched to new tissue culture dishes precoated with gelatin (from porcine skin, 0.1% g/v final, Sigma, G2500) without feeders in 2i/LIF medium [N2B27 basal medium (Neurobasal medium (50% v/v final, Gibco, 21103-049) and DMEM/F-12 medium (50% final, Gibco, 11320-074) supplemented with L-Glutamine (1.25 mM final, Gibco, 25030081), NDiff Neuro2 supplement (1x final, Millipore, SCM012), B27 supplement (1x final, Gibco, 17504-044), 0.8% (v/v) beta mercapto ethanol, and 1% (v/v) P/S) supplemented with 0.35% (g/v) Bovine Serum Albumin (BSA, Sigma, A7979), homemade mouse LIF, GSK3 inhibitor CHIR-99021 (3 μM final, Axon Medchem, Axon 1386) and MEK inhibitor PD0325901 (1 μM final, Axon Medchem, Axon 1408)] for four passages.

### Plasmids Constructs

The full-length mouse cDNAs of *Dusp9*, *Zic3*, *Dkc1, Otud6a, Fhl1, Zfp185*, and *Luciferase* (from pGL2-Basic Promage, E1641), *NLS-cherry* was cloned into pENTR vectors (Invitrogen, K240020) with either a C-terminal or a N-terminal HA tag, or no tag, and recombined into pPB-CAG-Dest-pA-pgk-bsd (PB-DEST-BSD) destination vectors. The PB-Scml2-BSD plasmid was obtained by recombining the pDONR221-Scml2 plasmid [56] into PB-DEST-BSD. Guide RNAs (gRNAs) were cloned into SapI digested pZB-sg3 [57]. All gRNAs sequences are included in S9 Table, S3 Fig F/H, S5 Fig A and S6 Fig B/E/H. All constructs were verified by DNA Sanger sequencing.

### Generation of stable male iPSCs overexpressing X-linked candidate genes

Male iPSCs (line 4, P5, grown on feeders in S/L conditions) were feeder-depleted before seeding in six-well plates precoated with 0.1% gelatin in S/L medium at a density of 650,000 cells per well, which were co-transfected with 1 ug of PB expression constructs encoding candidate genes and 3 ug of pCAGP Base [58] using 10 μl Lipofectamine 2000 (Invitrogen, 11668027). Transfected cells were selected with 20 μg/mL blasticidin (Fisher BioReagents, BP2647100) supplemented to the medium for two days starting from 24h after transfection and maintained with 5 μg/mL blasticidin thereafter.

### Generation of XX female ESC lines with heterozygous deletions of X-linked candidate genes

2000,000 female F1-2-1 ESCs (P19, grown on feeders in S/L condition) were resuspended in 1 ml of S/L medium and co-transfected with 2 ug of a plasmid expressing Cas9 under a CAG promoter and 1 ug of 2 plasmids (pZB-sg3 [57]) containing gRNAs (S9 Table) using 10 μl Lipofectamine 2000 (Invitrogen, 11668027) (S3 Fig F/H, S5 Fig A and S6 Fig B/E/H) for one hour before plating on 4-drug resistant (DR4) feeders. Transfected cells were selected with 2 μg/mL puromycin (Fisher BioReagents, BP2647100) on DR4 feeders in ESC medium for two days starting from 24h after transfection, and expanded at low density on WT feeders in 10cm dishes. Individual colonies were picked onto WT feeders, expanded for another two passages and genotyped for both WT and mutant alleles (primers in Table S8). WT and mutant alleles were further verified by DNA Sanger sequencing.

### Differentiation

To induce differentiation towards epiblast-like cells (EpiLCs), ESCs and iPSCs (male lines: 1, 4, 8, 12, 16; female lines: 19, 20, 21, 22, 23, 26; P8), which had been adapted to 2i/LIF conditions for 4 passages, were plated in N2B27 basal medium supplemented with 10 ng/ml Fibroblast Growth Factor-basic (Fgf2, Peprotech, 100-18C) and 20 ng/ml Activin A (ActA, Peprotech, 120-14E) on Fibronectin (5 ug/10 cm^2^, Millipore, FC010-5MG)-coated tissue culture plates at a cell density of 8*10^4^ cells/cm^2^ for four days, during which medium was refreshed daily and cells were harvested at different time points (0h, 12h, 1 day, 2 days, 3 days and 4 days), as previously described [15]. ESCs (WT female and male ESCs, *Dusp9+/-* ESCs, and *Zic3+/-* ESCs) and iPSCs (male iPSC lines 4, 8, 16; female iPSC lines 20, 21, 22; both early and late passages) grown in S/L condition were differentiated in the absence of feeders by LIF withdrawal (similar as mouse ESC medium but with 10% FBS and without LIF) at a cell density of 4*10^4^ cells/cm^2^ for two days, during which medium was refreshed daily and cells were harvested at different time points (0h, 24h and 48h), as previously described [15]. Likewise, male iPSC lines overexpressing X-linked genes were differentiated by LIF withdrawal with 5 μg/mL blasticidin in the absence of feeders.

### Clonal assays

iPSCs were subjected to LIF withdrawal, and 5000 cells were sorted onto feeders in S/L in each well of a 12-well plate in triplicate at 0h, 24h, 48h, 72h of LIF withdrawal. The next day cultures were switched to 2i/L. Alkaline Phosphatase (AP) staining was carried out 5 days after replating, using the VECTOR Red Alkaline Phosphatase (Red AP) Substrate Kit (VECTOR, SK-5100). Imaging was carried out using an Nikon Eclipse Ti2 Microscope equipped with an Nikon DS-Qi2 camera. AP positive colonies in each well of the 12-well plates were counted using NIS Element Auto Measurement.

### Cell growth assay

ESCs and iPSCs were plated in 24-well plates at a cell density of 4*10 ^4^ cells/cm^2^ for two days, during which medium was refreshed daily and cells were counted at different time points (0h, 12h, 24h, 36h and 48h). The cell numbers are presented as fold changes relative to cell numbers at 0h.

### FUCCI cell-cycle reporter assay

We generated XY and XX Mus/Cas ESCs expressing the FUCCI fluorescent reporters together with an H2B nuclear marker by co-transfecting of the WT XY and XX ESCs with PB expression constructs including PB-mCherry-hCdt1-BSD, PB-mVenus-hGeminin-PURO and PB-mCerulean-H2B-NEO [59] and pCAGP Base [58] using Lipofectamine 2000. Transfected cells were selected with 20 μg/mL blasticidin, 2 μg/mL puromycin and 100 μg/mL G418 supplemented to the medium for two days starting from 24h after transfection and maintained with 5 μg/mL blasticidin, 1 μg/mL puromycin and 50 ug/ml G418 thereafter. The BD FACSMelody cell sorter were used to analyze the FUCCI ESCs at different phases of the cell cycle.

### EdU incorporation assay

ESCs and iPSCs were pulse-labeled with the Click-iT EdU Alexa Fluor 647 Flow Cytometry Assay Kit (Invitrogen, C10424) according to the manufacturer’s instructions. Briefly, cells were incubated with 10μM 5-ethynyl-2′-deoxyuridine (EdU) for 45 min at 37°C. Then, cells were detached from plates with 0.05% Trypsin-EDTA (Gibco, 25300054), washed with PBS/ 2% BSA and aliquoted into one million cells per tube. Cells were fixed with 4% PFA for 20 minutes, washed with PBS/ 2% BSA and followed by 20 minutes permeabilization with PBS/ 0.5% Triton X-100. Cells were further incubated with the staining cocktail for 10 minutes at room temperature in the dark to reveal EdU incorporation. After twice washes with PBS/ 2% BSA, cells were stained with 3μM PI (Invitrogen, P1304MP) for 15 minutes at room temperature and analyzed using the BD FACSCanto II HTS flow cytometer.

### Immunofluorescence

Immunofluorescence analyses were carried out largely as described previously [27], using the following primary antibodies: NANOG (eBioscience, 14-5761 clone eBioMLC-51, 1/200; and Abcam, ab80892, 1/ 200), DPPA4 (R&D, AF3730, 1/200), HA (Cell Signaling Technology, 2367S, 1/100), DUSP9 (Abcam, ab167080, 1/100). Images were acquired using an ApoTome Zeiss Microscope equipped with an AxioCam MRc5 camera. ESC and iPSC lines were defined as NANOG+ or DPPA4+ when >50% cells showed NANOG or DPPA4 staining signal.

### RNA FISH

RNA Fluorescence In Situ Hybridization (RNA FISH) analyses were carried out mostly as described previously using double stranded directly labelled DNA probe for *Tsix/Xist [27]*. Images were acquired using an ApoTome Zeiss Microscope equipped with an AxioCam MRc5 camera. Single-cell resolution analysis of *Tsix/Xist* biallelic expression in iPSCs and ESCs was determined by calculating the ratio of cells with biallelic *Tsix/Xist* expression to the cells with monoallelic or biallelic *Tsix/Xist* expression.

### Genomic DNA extraction and qPCR

Genomic DNA (gDNA) was extracted from feeder-depleted ESCs and iPSCs using the PureLink Genomic DNA Kit (Invitrogen, K1820) and qPCR was performed using the Platinium SYBR Green qPCR SuperMix-UDG kit (Invitrogen, 11733046) on a ABI ViiA7 real-time PCR system (Applied Biosystems), following the manufacturer’s protocol. Primers against four X-linked genes (*Tfe3, Bcor, Pdha1, and Mid1*) covering the two distal parts of the mouse X-chromosome are listed in S9 Table (Fig 1 G). The standard curve was derived from serial dilutions of gDNA from XY ESCs (V6.5). All qPCR assays used had an efficiency above 95%. Relative quantities of each gene were measured as arbitrary units from comparison to the standard curve. The ratio of X-chromosome to autosome (X/Autosome Ratio) in DNA level was presented as the average ratio of the X-linked gene quantity (*Tfe3, Bcor, Pdha1* and *Mid1*) to the autosomal gene quantity (*Gapdh*), in other words X/Autosome Ratio = (*Tfe3/Gapdh* + *Bcor/Gapdh* + *Pdha1/Gapdh* + *Mid1/Gapdh*)/4.

### RT-qPCR

Total RNA was extracted using the RNeasy Mini Kit (Qiagen, 74106) or TRIzol (Invitrogen, 15596026). cDNA synthesis was performed using the SuperScript III First-Strand Synthesis SuperMix kit (Invitrogen, 11752-050) and RT-qPCR was performed using the Platinium SYBR Green qPCR SuperMix-UDG kit (Invitrogen, 11733046) and on the ABI ViiA7 real-time PCR system, following the manufacturer’s protocol. Primers used are listed in S9 Table. The standard curve was derived from serial dilutions of cDNA. All assays used had an efficiency above 95%. Relative quantities of each transcript were calculated as arbitrary units from comparison to the standard curve. Relative expression level of the target transcript was presented as the ratio of the target transcript quantity to the housekeeping transcript (Gapdh) quantity. Logarithm values (base 2) of relative expression levels were used for assessment of the gene expression kinetics during differentiation. The relative gene expression levels of five pluripotency-associated genes (*Prdm14, Nanog, Tcl1, Rex1* and *Esrrb*) from iPSCs (male lines: 1, 4, 8, 12, 16; female lines: 19, 20, 21, 22, 23, 26) and ESCs (V6.5 male ESCs and F1-2-1 female ESCs) at 0h and 24h of EpiLC differentiation were used for unsupervised clustering comparison, which was performed in R with heatmap.2 function in package “gplots”.

### RNA sequencing

Total RNA was isolated from two independent female *Dusp9*+/- ESC lines, *Dusp9+/+* XX and XY ESCs in both the undifferentiated state and the differentiated state after 24 hours of LIF withdrawal using TRIzol following the manufacturer’s protocol. 4 μg of total RNA was used for construction of stranded poly(A) mRNA-Seq library with the KAPA stranded mRNA Library prep kit (KAPA Biosystems, KK8421). Library concentrations were quantified with the Qubit dsDNA HS (High Sensitivity) Assay Kit (Invitrogen, Q32854), and equimolar amounts were pooled for single-end sequencing on an Illumina HiSeq 4000 instrument (Illumina) to yield ~20 million (range 16-23 million) 36bp long reads per sample.

### Differential gene expression analysis

Reads from all datasets (*Dusp9+/-* ESCs, *Dusp9+/+* ESCs and XY ESCs) were aligned to mouse reference genome GRCm38/mm10 using STAR (v2.5.3a) with default parameters followed by conversion to BAM format sorted by coordinate. The mapping efficiencies of the datasets were >69% of uniquely mapped reads. Subsequently, the featureCounts function from the R Bioconductor package “Rsubread” was used to assign mapped reads to genomic features. For downstream analyses, only the genes with CPM value (count-per-million) higher than 0.5 in at least two libraries were retained. The resulting read count matrix (S8 Table) was used as the input for PCA with the top 500 most variable genes. Differential gene expression analysis was performed using the edgeR quasi-likelihood pipeline in R [60]. Obtained p-values were corrected for multiple testing with the Benjamini-Hochberg method to control the FDR. DEGs were defined on the basis of both FDR < 0.05 and fold difference ≧ 1.5. Venn diagrams were generated using an online tool as previously described [61]. Heatmaps were created using unsupervised hierarchical clustering of both 200 most variable genes and the different samples and generated in R using the heatmap.2 function of the package “gplots”.

### Omni-ATAC-seq

Assay for transposase accessible chromatin (ATAC) followed by sequencing was performed using the Omni-ATAC protocol [34]. Briefly, iPSCs and ESCs were expanded on top of male WT feeders in mouse ESC medium (S/L condition). After feeder-depletion, 50,000 viable cells were pelleted at 500 RCF at 4°C for 5 min in a fixed angle centrifuge, and then the cells were gently washed once with 50 μl of cold PBS. Next, the cell pellets were resuspended in 50 μl of ATAC-lysis buffer (10mM Tris HCl pH7.4, 10mM NaCl, 3mM MgCl2, 0.1% Tween-20, 0.1% NP40, and 0.01% Digitonin) and incubated on ice for 3 min. Wash out lysis with 1 ml of cold ATAC-lysis buffer containing 0.1% Tween-20 but No NP40 or digitonin and invert tube 3 times to mix. Nuclei were pelleted at 500 RCF for 10 min at 4°C in a fixed angle centrifuge. After discarding all supernatant, nuclei were resuspended in 50 μl of transposition mixture (25 ul 2x TD buffer, 2.5 ul transposase (100 nM final), 16.5 ul PBS, 0.5 ul 1% digitonin, 0.5 ul 10% Tween-20, and 5 ul H2O) (Nextera DNA Sample Preparation Kit, Illumina, FC-121-1030). The reaction was performed at 37°C for 30 minutes in a thermomixer with 1000 RPM mixing. The transposed DNA was purified using a Zymo DNA Clean and Concentrator-5 Kit (D4014). DNA libraries were PCR amplified using NEBNext High-Fidelity 2x PCR Master Mix (Bioke, M0541), and size selected for 200 to 800 bp using homemade Serapure beads [62]. Library concentrations were quantified with the KAPA Library Quantification Kit (KK4854), and equimolar amounts were pooled for single-end sequencing on an Illumina HiSeq 4000 instrument (Illumina) to yield ~50 million (range 34-90 million) 51bp long reads per sample.

### Differential chromatin accessibility analysis

Single-end ATAC-seq raw data were analyzed using the ATAC-seq pipeline from the Kundaje lab (Version 0.3.3) [63]. Briefly, the raw reads were first trimmed using cutadapt (version 1.9.1) to remove adaptor sequence at the 3′ end. The trimmed reads were aligned to reference genome (mm10) using Bowtie2 (v2.2.6) using the ‘--local’ parameter. Single-end reads that aligned to the genome with mapping quality ≥30 were kept as usable reads (reads aligned to the mitochondrial genome were removed) using SAMtools (v1.2). PCR duplicates were removed using Picard’s MarkDuplicates (Picard v1.126). Open chromatin regions (peak regions) were called using MACS2 (v2.1.0) using the ‘-g 1.87e9 -p 0.01 --nomodel --shift - 75 --extsize 150 -B --SPMR --keep-dup all --call-summits’ parameter [64]. The differential chromatin accessibility analysis and related plots were performed using the DiffBind package with ‘DESeq2, log2fold=1, FDR<=0.05’ parameter [65]. GO analysis for Biological Process terms was performed using GREAT (v3.0.0) analysis [66] with the mm10 reference genome, where each region was assigned to the single nearest gene within 1000 kb maximum distance to the gene’s TSS.

### Motif Discovery Analysis

Known motif search was performed using program of findMotifsGenome.pl in the HOMER package (v4.9.1) with ‘mm10 -size -250,250 -S 15 -len 6,8,10,12,16’ parameters [67]. Incidences of specific motif was examined by the program of annotate-Peaks.pl in the HOMER package with size parameter ‘‘-size 500’’.

### Western blots

Cells were detached from plates with 0.25% Trypsin-EDTA (Gibco, 25200056), pelleted before addition of RIPA lysis buffer (Sigma, R0278-50ML) supplemented with 1% (v/v) Protease inhibitor cocktail (Sigma, P8340-1ml) and 1% (v/v) Phosphatase inhibitor Cocktail 3 (Sigma, P0044-1ML), and lysed on ice for 30 min. The lysates were spun for 10 min at 13000 rpm. The protein concentration was determined with BCA protein assay kit (Pierce, 23225). Each sample with 15 μg of total protein was denatured in 1x LDS Sample buffer (Life Technologies, NP0007) with 100 mM DTT for 5 min at 98°C. The cell lysates were loaded onto a 4%–15% mini-Protean TGX gel (Bio-Rad, 456-1083), electrophoresed, and transferred to nitrocellulose membranes (VWR,10600002). Membranes were blocked in PBS 0.1% (v/v) Tween-20 and 5% (g/v) blotting reagent (Bio-Rad, 1706404) and incubated with the following primary antibodies overnight at 4°C: rabbit anti-NANOG (Abcam, ab80892, 1/1000), rabbit anti-DUSP9 (Abcam, ab167080, 1/500), mouse anti-DKC1 (Santa Cruz, sc-365731, 1/250), mouse anti-HA (Cell Signaling Technology (CST), 2367S, 1/1000), sheep anti-ZIC3 (R&D Systems, AF5310, 1/250) and mouse anti-ACTIN (Abcam, ab3280, 1/5000). After extensive PBS 0.1% Tween-20 (PBS-T) washes, membranes were incubated with a secondary HRP-conjugated goat anti-mouse IgG antibody (Bio-Rad, 1706516, 1/5000) or goat anti-rabbit IgG antibody (Bio-Rad, 1706515 1/5000) for 30 minutes at room temperature. After another round of extensive PBS-T washes, protein expression was visualized using the ECL chemiluminescence reagent (Perkin-Elmer, NEL103001EA) and LAS-3000 imaging system (Fuji). Data were analyzed with ImageJ.

### Statistical analysis

Statistical tests were performed using the Graphpad Prism 5 software (GraphPad Software). Unpaired two-tailed t-test, one-way ANOVA with multiple comparisons test or two-way repeated-measures ANOVA were used as indicated. All data are presented as the mean±SEM. p-values <0.05 were considered statistically significant.

### Single Cell RNA-seq analysis

Single cell RNA-seq data of XX female and XY male ESCs in S/L and 2i/L from published data set [24] was re-aligned to N-masked mouse reference genome mm10 using hisat 2 (2.0.5) with disabled soft-clipping. Alignment was followed by conversion to BAM files using SAMtools 1.4.1, aligned reads were then summarized using featureCounts v1.5.2. Quality controls and downstream analyses were performed with the use of scater and SingleCellExperiment packages [68,69]. Cells displaying total counts lower than 500,000 reads and less than 9000 genes detected were discarded from the analysis.

### Data Availability

The GEO accession number for the RNA-seq and ATAC-seq data reported in this paper is GSE110215. The single cell RNA-seq data from published data [24] was deposited under the accession number GSE74155 NCBI Gene Expression Omnibus database.

## ACKNOWLEDGEMENTS

We apologize to the authors that we could not cite due to space constraint. We thank Stein Aerts, Konrad Hochedlinger, Frederic Lluis Vinas and Edda Schulz for discussions; Rudolf Jaenisch for providing mice; Qiaolin Deng for providing Mus/Cas ESCs, Rita Khoueiry and Michela Bartoccetti for help with derivation of Mus/Cas ESCs, Ye-Guang Chen for the *Dusp9* plasmids, Miguel Branco for the Scml2 plasmids, José Silva for the PiggyBac plasmid, Mitchell Guttman and Jesse Engreitz for the pZB-sg3 plasmid, Ali Brivanlou and Ariel Waisman for the Fucci plasmids, Stein Aerts, Kristofer Davie, Liesbeth Minnoye and Xinlong Luo for help with bioinformatics and ATAC-seq; Kenjiro Shirane and Hiroyuki Sasaki for providing processed data. We thank Constantinos Chronis, Kian Koh, Kathrin Plath and Edda Schulz for comments on the manuscript. We are grateful to the help of the KU Leuven FACS Core, Genomics Core and Mouse facility, Metabolomics Core, VIB/KU Leuven and SCIL.

## AUTHOR CONTRIBUTIONS

Conception and design, J.S. and V.P.; Experiments, J.S., N.D.G., L.V., T.O., I.T. and V.P.; Analyses, J.S; ATAC-seq and RNA-seq analyses, A.J.and J.S.; Writing V.P. and J.S. with input from all authors; Supervision, V.P.

## SUPPLEMENTAL FIGURES AND LEGENDS

S1 Fig. X-dosage-specific differences in transcriptome, pluripotency exit and cell growth in mouse iPSCs.

**(A)** Immunofluorescence analysis for NANOG and DPPA4 in iPSCs and ESCs grown in 2i/L. MEFs served as negative control. Representative images for NANOG (Red), DPPA4 (Green) and Dapi (Blue, nuclei counterstaining) are shown.

**(B)** Summary of ESC and iPSC lines used in this study as well as NANOG and DPPA4 protein expression analysis in 2i/L and in S/L.

**(C)** RNA FISH analysis for *Tsix/Xist* expression in iPSCs grown in 2i/L. Representative images for *Tsix/Xist* RNA (Green) and Dapi (Blue, nuclei counterstaining) are shown. Yellow arrowheads point to *Tsix/Xist* transcriptional sites. Also see [1]. Right: Quantification of *Tsix/Xist* RNA FISH signal from Figure S1C and for all lines, plotted as the proportion of cells with biallelic *Tsix/Xist* signal (= number of cells with biallelic *Tsix/Xist* expression / number of cells with biallelic or monoallelic expression). The number of counted nuclei is > 50 per cell line.

**(D)** qPCR analysis for X-chromosome DNA copy number in both early passage and late passage iPSCs grown in S/L. X copy number are presented as the average ratio of gDNA quantities for four X-linked genes (*Tfe3, Bcor, Pdha1, and Mid1*, locations in X-chromosome shown in lower panel) to gDNA quantities for autosomal gene *Gapdh*.

**(E)** Unsupervised hierarchical clustering of stem-cell maintenance gene expression in XX, XY and XO iPSCs.

**(F)** RT-qPCR analysis for pluripotency-associated gene expression in XX, XY and XO iPSCs grown in S/L. The expression values are represented as averages (±SEM) of XY, XX and XO iPSC lines (three different lines each) in early passage (P8) and late passage (P14), respectively. Statistical significance was analysed using the unpaired, two-tailed t-test (**p<0.01).

**(G)** DEG analysis, identifying clear differences between XX and XY ESCs.

**(H)** Venn diagrams showing the overlap between XX vs XY iPSCs DEGs and XX vs XY ESCs DEGs. Despite difference in genetic background between iPSCs and ESCs, 93 genes are differentially expressed between XX and XY cells both for iPSCs and ESCs.

**(I)** RT-qPCR for pluripotency-associated genes for individual XY (blue), XX (magenta) and pXO (orange, line 26) iPSC lines undergoing EpiLC differentiation.

**(J)** Unsupervised hierarchical clustering of XX, XY and XO iPSC and ESC lines based on expression of X-linked genes in undifferentiated state (i), and pluripotency-associated genes in undifferentiated state (ii) and 24h of LIF withdrawal differentiated state (iii).

**(K)** Growth curves and doubling times of XY and XX iPSCs (i) and ESCs (ii) in 2i/LIF condition. Cell counts were obtained at different time points, as indicated. Cell counts are shown as fold changes relative to 0h and represent the averages cell count (±SEM) over three cell lines for male and female iPSC lines (n=1, left panel) at early passage (P8) and for two male and female ESCs lines (n=1, right panel). Growth curve: *p* value <0.001 (***) between male and female cell lines, by two-way repeated-measures ANOVA with Bonferroni posttests. Td: \**p*<0.05, male lines vs female lines, by unpaired two-tailed t-test.

**(L)** (i) EdU incorporation in combination with DNA staining using PI shows a higher number of XX iPSCs residing in S phase, and a lower number of XX iPSCs residing in G1 phase when compared to XY iPSCs. Results show the averages (±SEM) over three XX female iPSC lines and three XY male iPSC lines (n=1). Significance was tested using one-way ANOVA with Sidak’s multiple comparisons test (**p<0.01, ***p<0.001). (ii) FUCCI reporter expression shows the number of XX female ESCs activating the G1 phase reporter is lower than that of XY male ESCs, and the number of XX female ESCs activating the S/G2/M phase reporter is higher than that of XY male ESCs. Results show the averages (±SEM) over two XX female and two XY male FUCCI reporter ESC lines (n=1). Significance was tested using one-way ANOVA with Bonferroni’s multiple comparisons test (**p<0.01, ***p<0.001).

**(L)** XY and XX ESCs competition assay. WT XX ESCs were mixed with GFP-labelled XY ESCs (Oct4-GiP) in different ratios (left panel, scheme of the experiment) and the proportion of GFP-labeled cells in the culture was measured over time (n=2), as indicated. *p<0.05, **p <0.01, by two-way repeated-measures ANOVA with Dunnett’s multiple comparisons test compared to day 0.

S2 Fig. Influence of X-dosage on the chromatin regulatory landscape of mouse iPSCs and ESCs.

**(A)** Distance to closest TSSs of “Not differentially accessible”, “XX gain” and “XY gain” ATAC-seq regions in iPSCs.

**(B)** Mean read count ratio to autosomes showing which lines have increased/decreased ATAC-seq reads on the Y chromosome, X chromosome, chromosome 8 and chromosome 9. This analysis confirms the higher abundance of DNA sequence reads coming from the Y chromosome in XY lines, and from the X chromosome in XX lines.

**(C)** Sample-to-sample distance heatmap showing the Euclidean distances (calculated from the rlog transformed counts, DESeq2) between ESCs samples. Samples cluster by X-dosage (XX vs XY).

**(D)** Differential chromatin accessibility analysis between XX and XY ESCs. Log2 fold change (XX/XY) in reads per accessible region are plotted against the mean reads per ATAC-seq peak. Thousands of open chromatin regions that more open in XX ESCs or in XY ESCs (|log2fold|>=1, false discovery rate (FDR)<=0.05).

**(E)** Venn diagrams showing the overlap between genes nearest to the “XX gain” or "XY gain" regions defined in (D) and the DEGs between XX and XY ESCs (DEGs= |log_2_fold|>=log_2_1.5, FDR<=0.05).

**(F)** Transcription factor motifs enriched in chromatin regions more open in XX ESCs.

**(G)** Transcription factor motifs enriched in chromatin regions more open in XY ESCs.

**(H)** Overview of motif enrichment in XX gain and XY gain chromatin regions in ESCs.

**(I)** Overview of motif enrichment in open chromatin not differentially accessbilie between XX and XY cells in both ESCs and iPSCs.

**(J)** Transcription factor motifs enriched in open chromatin not differentially accessbilie between XX and XY iPSCs.

S3 Fig. Effects of Zic3 overexpression and Zic3/ Tfe3 heterozygous deletion in XX ESCs on pluripotency exit.

**(A)** Expression of *Zic3* in XY, XX and XO iPSCs showing 2.1 fold increased *Zic3* dosage in XX iPSCs over XY and XO iPSCs as assessed by RNA-seq.

**(B)** Scheme of *Zic3* overexpression in XY iPSCs, followed by characterization and LIF withdrawal differentiation.

**(C)** (i) Western blot analysis for ZIC3 and ACTIN in XY iPSCs after HA-Zic3 or Zic3-HA overexpression. (ii) quantification using ACTIN as loading control. ZIC3 protein values are represented as averages (±SEM) of three independent experiments.

**(D)** RT-qPCR of *Zic3* in HA-tagged Zic3 overexpressing (3.3-3.6 fold) XY iPSCs. XY iPSCs overexpressing Luciferase served as negative control. Results are presented as averages (±SEM) of three independent experiments.

**(E)** RT-qPCR analysis for *Zic3*, *Prdm14*, *Nanog* and *Tcl1* expression during LIF withdrawal in XY iPSCs overexpressing HA-Zic3, Zic3-HA or Luciferase control. Results are presented as averages (±SEM) of three independent experiments (n=3).

**(F)** Scheme of heterozygous *Zic3* deletion strategy in XX ESCs. The sequences of the gRNAs used to deleted *Zic3* are shown in blue with PAM sequences in red. Two independent *Zic3*+/- XX ESC lines were derived. The sequences of the knockout (KO) alleles were obtained by Sanger sequencing. Red line shows the location of the gRNAs. Green arrows shows the locations of the primers for genotyping PCR.

**(G)** Genotyping of *Zic3* heterozygous deleted XX ESC lines for both WT and KO alleles. The parental *Zic3+/+* ESCs were used as positive control for the WT allele and as negative control for the KO allele.

**(H)** as in (F) but for heterozygous *Tfe3* deletion strategy in XX ESCs.

**(I)** Genotyping of *Tfe3* heterozygous deleted XX ESC lines for both WT and KO alleles.

**(J)** qPCR analysis for X-chromosome DNA copy number. X copy number are presented as the relative gDNA quantities for six X-linked genes (*Tfe3, Bcor, Nr0b1, Otud6a, Pdha1, and Mid1*, locations in X-chromosome shown in Figure S1H) to gDNA quantities for autosomal gene *Gapdh*.

S4 Fig. Overexpression of X-linked candidates Dkc1, Otud6a, Fhl1, Zfp185 and Scml2 has no effect on the pluripotency exit kinetics of XY iPSCs.

**(A)** Scheme for candidate X-linked gene overexpression in XY iPSCs, followed by LIF withdrawal.

**(B)** Map of the X-chromosome showing candidate X-linked genes.

**(C)** Western blot analysis for HA-tagged DKC1, OTUD6A, FHL1, ZFP185 using an anti-HA antibody. ACTIN was used as a loading control. Representative images are shown.

**(D)** Immunofluorescence analysis for NANOG and HA in stable XY iPSC lines overexpressing HA-tagged *Dkc1*. The stable XY iPSC line overexpressing Luciferase was used as a negative control. Representative images for NANOG (Red), HA (Green) and Dapi (Blue, nuclei counterstaining) are shown.

**(E-I) (i)** Expression of X-linked candidates *Dkc1, Otud6a, Fhl1, Zfp185 and Scml2* in XY, XX and XO iPSCs showing fold change (fc) in XX iPSCs over XY iPSCs as assessed by RNA-seq. **(ii)** RT-qPCR of X-linked candidates expression in XY iPSC lines with ectopic expression of HA-tagged *Dkc1* (E), HA-tagged *Otud6a* (F), HA-tagged *Fhl1* (G), HA-tagged *Zfp185* (H) or *Scml2* (I). XY iPSCs overexpressing Luciferase served as negative control. **(iii)** RT-qPCR for pluripotency-associated genes *Prdm14*, *Nanog* and *Tcl1*, and the respective X-linked candidate genes in XY iPSC lines stably overexpressing the respective HA-tagged X-linked candidates and subjected to LIF withdrawal. Results are presented as averages (±SEM) of three (E) or two (F-J) independent experiments, which are not statistically significant by two-way repeated-measures ANOVA with Bonferroni posttests.

S5 Fig. Characterization of Dusp9 heterozygous mutant XX female ESCs

**(A)** Scheme of heterozygous *Dusp9* deletion in XX ESCs. The gRNAs sequences used to delete the *Dusp9* gene are shown in blue with PAM sequences in red. Two independent *Dusp9+/-* ESC lines were derived. The sequences of the KO alleles were obtained by DNA Sanger sequencing. Red line shows the location of the gRNAs. Green arrows shows the locations of the primers for genotyping PCR.

**(B)** Genotyping *Dusp9*+/- ESC lines for both the WT and the KO allele. The parental XX ESC line was used as a positive control for the WT allele PCR and as a negative control for the KO allele PCR.

**(C)** RNA FISH analysis for *Tsix/Xist* expression in the two independent *Dusp9+/-* ESC lines and their parental XX ESC line. Representative images for *Tsix/Xist* RNA (Green) and Dapi (Blue, nuclei counterstaining) are shown. Yellow arrowheads point to *Tsix/Xist* transcriptional sites. Right: Quantification of Figure S5C, plotted as the proportion of cells with biallelic *Tsix/Xist* signal (= number of cells with biallelic *Tsix/Xist* expression / number of cells with biallelic or monoallelic expression). Numbers of counted nuclei > 50 per cell line.

**(D)** (i) qPCR analysis for X-chromosome DNA copy number. X copy number are presented as the average ratio of gDNA quantities for four X-linked genes (Tfe3, Bcor, Pdha1, and Mid1) to gDNA quantities for autosomal gene Gapdh. Results are presented as averages (±SEM) of the same lines in two independent qPCR experiments (n=2). (ii) Mean read count ratio to autosomes showing which lines have increased/decreased ATAC-seq reads on the Y chromosome, X chromosome, chromosome 8 and chromosome 9. This analysis confirms the higher abundance of DNA sequence reads coming from the Y chromosome in XY lines, and from the X chromosome in XX lines. (iii) Mean expression ratio to autosomes for sex chromosomes and chromosomes 8 and 9.

**(E)** DNA methylation analysis of *Dusp9*+/- ESCs, *Dusp9*+/+ ESCs and XY ESCs by mass spectrometry. (i) 5mC, (ii) 5hmC.

**(F)** RNA-seq analysis of pluripotency associated gene expression during pluripotency exit in Dusp9+/- and Dusp9+/+ ESCs. The delay in pluripotency gene downregulation is maintained.

**(G)** Scheme of *Dusp9* overexpression in male iPSCs grown in S/L, followed by characterization and LIF withdrawal.

**(H)** (i) Western blot analysis for DUSP9 in XY iPSC lines with ectopic HA-tagged DUSP9 expression. ACTIN was used as a loading control. (ii) quantification using ACTIN as loading control. DUSP9 protein values are represented as averages (±SEM) of three independent experiments (n=3).

**(I)** Immunofluorescence analysis for NANOG and HA-tagged DUSP9 in stable male iPSCs with HA-tagged DUSP9 overexpression. The stable male iPSC line overexpressing Luciferase was used as a negative control. Representative images for NANOG (Red), HA (Green) and Dapi (Blue, nuclei counterstaining) are shown.

**(J)** Expression of *Dusp9* in XX, XY and XO iPSCs showing 2.7 fold increased *Dusp9* dosage in XX iPSCs over XY and XO iPSCs as assessed by RNA-seq.

**(K)** qRT-PCR for *Dusp9* in XY iPSCs overexpressing HA-Dusp9, Dusp9-HA and control Luciferase.

**(L)** qRT-PCR for *Dusp9* and pluripotency-associated genes *Prdm14*, *Nanog* and *Tcl1* in XY iPSCs overexpressing *Dusp9* following LIF withdrawal. The stable XY iPSC line overexpressing Luciferase served as a negative control. Results are presented as averages (±SEM) of three independent experiments.

S6 Fig. Large fragment heterozygous deletion of X chromosome in XX ESCs.

**(A)** Scheme of the three large fragment (LF) heterozygous deletions in XX female ESCs.

**(B)** Scheme of heterozygous LF1 deletion in XX ESCs. The gRNAs sequences are shown in blue with PAM sequences in red. Two independent *LF1*+/- ESC lines were derived. The sequences of the KO alleles were obtained by DNA Sanger sequencing. Red line shows the location of the gRNAs. Green arrows shows the locations of the primers for genotyping PCR used in (C).

**(C)** Genotyping *LF1*+/- XX ESC lines for both the WT and the KO allele.

**(D)** qPCR analysis for X-chromosome DNA copy number. X copy number are presented as the relative gDNA quantities for six X-linked genes (Tfe3, Bcor, Pdha1, and Mid1) to gDNA quantities for autosomal gene Gapdh. Results are presented as averages (±SEM) of the same lines in two independent qPCR experiments (n=2).

**(E-G)** As in (B-D) for heterozygous LF2 deletion in XX ESCs. Two independent *LF2*+/- ESC lines were derived and validated.

**(H-J)** As in (B-D) for heterozygous LF3 deletion in XX ESCs. Three independent *LF3*+/- ESC lines were derived and validated.

S7 Fig. Single cell RNA-seq analysis of XX female and XY male ESCs in S/L and 2i/L.

**(A)** PCA analysis of XX female and male XY ESCs S/L and 2i/L single cell RNA-seq data from [2]. Cells grown in S/L are shown in magenta, those grown in 2i/L are shown in blue.

**(B)** Same analysis as in (A), showing XX female ESCs in magenta and XY male ESCs in blue.

**(C)** Expression of *Nanog*, *Prdm14*, *Esrrb* and *Tcl1* projected onto the PCA shown in (A) and (B).

**(D)** Expression (log read count) of pluripotency associated genes *Nanog*, *Prdm14*, *Tcl1*, *Esrrb* and *Zfp42* in single XX female and XY male ESCs grown in S/L. Violin plots indicate the distribution of single cells where each dot is a cell. Black lines indicate median gene expression.

**(E)** As in (D) for the same cells grown in 2i/L.

**SUPPLEMENTAL TABLES**

S1 Table. Summary of cell lines used in this study.

S2 Table. Differentially expressed genes between XX and XY iPSCs.

S3 Table. Differentially expressed genes between XX and XY ESCs.

S4 Table. Differential open chromatin regions in XX and XY iPSCs.

S5 Table. Differential open chromatin regions in XX and XY ESCs.

S6 Table. Differentially accessible regions associated with differential gene expression.

S7 Table. Expression level of X-linked candidate genes in XX and XY iPSCs and ESCs.

S8 Table. RNA-seq data, CMP counts.

S9 Table. Primer sequences.

